# The striatal heterogeneous nuclear ribonucleoprotein H mRNA targetome associated with methamphetamine administration and behavior

**DOI:** 10.1101/2021.07.06.451358

**Authors:** Qiu T. Ruan, William B. Lynch, Rebecca H. Cole, Michael A. Rieger, Britahny M. Baskin, Sophia A. Miracle, Jacob A. Beierle, Emily J. Yao, Jiayi W. Cox, Amarpreet Kandola, Kayla T. Richardson, Melanie M. Chen, Julia C. Kelliher, R. Keith Babbs, Peter E. A. Ash, Benjamin Wolozin, Karen K. Szumlinski, W. Evan Johnson, Joseph D. Dougherty, Camron D. Bryant

## Abstract

Methamphetamine addiction remains a major public health concern in the United States that has paralleled the opioid epidemic. Psychostimulant use disorders have a heritable genetic component that remains unexplained. Methamphetamine targets membrane and vesicular transporters to increase synaptic dopamine, norepinephrine, and serotonin. We previously identified *Hnrnph1* (heterogeneous nuclear ribonucleoprotein H1) as a quantitative trait gene underlying methamphetamine behavioral sensitivity. *Hnrnph1* encodes the RNA-binding protein hnRNP H1 that is ubiquitously expressed in neurons throughout the brain. Gene-edited mice with a heterozygous frameshift deletion in *Hnrnph1’s* first coding exon of showed reduced methamphetamine-induced dopamine release and behaviors. To inform the mechanism linking hnRNP H with methamphetamine neurobehavioral effects, we surveyed the mRNA targetome of hnRNP H via cross-linking immunoprecipitation coupled with RNA-sequencing in striatal tissue at baseline and at 30 min post-methamphetamine in wild-type male and female C57BL/6J mice. Methamphetamine induced changes in RNA-binding targets of hnRNP H in mice, including differential binding to 3’UTR targets and multiple enriched mRNAs involved in synaptic plasticity. Targetome, transcriptome, and spliceome analyses triangulated on *Cacna2d2* as a suggestive target, with differences in hnRNP H binding, gene expression and splicing following methamphetamine treatment (2 mg/kg, i.p.). Furthermore, pre-treatment with pregabalin, an inhibitor of α2δ2 and α2δ1 voltage-gated calcium channel subunits, attenuated methamphetamine-induced locomotor activity in male and female mice, supporting a role for Cacna2d1/d2 in methamphetamine locomotor stimulant sensitivity. Our study identifies a dynamic hnRNP H RNA targetome that can rapidly and adaptively respond to methamphetamine to regulate gene expression and likely synaptic plasticity and behavior.

## 1. INTRODUCTION

Methamphetamine has significant misuse liability and induces neurotoxicity (Kish, 2008; Galbraith, 2015). In the U.S., drug overdose deaths involving psychostimulants increased ∼8-10-fold from 2012-2022 and continue rising (Spencer, 2024). There are no FDA-approved treatments for psychostimulant disorder which includes both methamphetamine and cocaine. Methamphetamine is a substrate for membrane dopamine transporters (and norepinephrine and serotonin) and vesicular monoamine transporters and causes reverse transport of dopamine from the presynaptic terminal and increased dopamine release (Fleckenstein et al., 2009; Siciliano et al., 2014). Increased dopamine release is a major contributor to methamphetamine’s misuse liability (Baumann et al., 2002). The rapid cell biological adaptations following methamphetamine action are not fully understood but could inform treatments for restoring cellular/synaptic function.

RNA binding proteins (**RBPs**) such as hnRNP H1 (coded by *Hnrnph1*; heterogeneous nuclear ribonucleoprotein H1; codes for hnRNP H1) regulate all aspects of RNA metabolism (Darnell, 2013; Hentze et al., 2018). RBP-coding genes are implicated in multiple substance use disorders, yet their dynamic role in regulating drug-induced changes in gene expression remains unexplored (Bryant and Yazdani, 2016). For example, deletion of fragile X mental retardation protein (**FMRP**) disrupts dopaminergic neuron development and alters psychostimulant-induced neuroplasticity and behavior (Fulks et al., 2010; Fish et al., 2013; Smith et al., 2014; Huebschman et al., 2021). Furthermore, alternative splicing of exon 2 and transcriptional regulation of *Oprm1* (codes for mu opioid receptor) are mediated through binding of hnRNP H1 and recruitment of other hnRNPs to its intronic AGGG sequence (Xu et al., 2014), suggesting hnRNPs can regulate transcripts of direct molecular targets of misused substances. Interestingly, coding mutations in *HNRNPH1* and *HNRNPH2* lead to severe neurodevelopmental disorders (Pilch et al., 2018; Reichert et al., 2020).

We previously mapped and validated *Hnrnph1* as a quantitative trait gene for methamphetamine stimulant sensitivity in mice (Yazdani et al., 2015) and identified a set of four 5’UTR variants that were functionally associated with decreased hnRNP H protein and methamphetamine-induced behavior (Ruan et al., 2020b). We also generated mice heterozygous for a 16 bp indel within the first coding exon of *Hnrnph1* (“*Hnrnph1* mutants”) that also showed reduced methamphetamine-induced locomotion, reward, reinforcement, and ventral striatal dopamine release (Ruan et al., 2020a). Striatal, synaptosomal proteomics identified opposite methamphetamine-induced changes in mitochondrial protein expression between genotypes, suggesting mitochondrial perturbations could underlie reduced methamphetamine-induced dopamine release and behavior in the mutants (Ruan et al., 2020a).

*Hnrnph1* codes for an RBP expressed throughout the brain that belongs to the hnRNP H/F subfamily of hnRNP RBPs that engage in all aspects of RNA processing (Honoré et al., 1995; Arhin et al., 2002; Han et al., 2010; Geuens et al., 2016; Uren et al., 2016). During cellular stress, RBPs (Markmiller et al., 2018) including hnRNP H (Wall et al., 2020), hnRNP A1 (Guil et al., 2006), and hnRNP K (Fukuda et al., 2009) localize to cytoplasmic stress granules and sequester mRNAs from translation. These hnRNP H-associated stress granules suggest hnRNP H interactions with its target mRNAs can change rapidly following perturbations in the cellular environment.

Methamphetamine-induced dopamine release within the nucleus accumbens contributes to its stimulant and addictive properties (Wise, 2004). Because we observed reduced methamphetamine-induced ventral striatal dopamine release and behaviors in *Hnrnph1* mutants (Yazdani et al., 2015; Ruan et al., 2020a), here, we sought to identify the striatal RNA-binding targets of hnRNP H in wild-type mice. We examined RNA-binding, mRNA expression, and splicing in *Hnrnph1* at baseline and following methamphetamine administration. We hypothesized that methamphetamine would alter RNA-binding, splicing, and expression of hnRNP H-bound genes. hnRNP H is primarily localized to the nucleus but can also be detected in cytoplasm (Wall et al., 2020). Within the striatal tissue, we predicted we would identify both presynaptic (e.g., dopaminergic terminals originating from midbrain cell bodies) and post-synaptic (striatal neurons) RNA targets relevant to dopaminergic transmission and signaling that could link hnRNP H to decreased methamphetamine-induced dopamine release and behavior. Our results triangulated on Cacna2d2 (voltage-dependent calcium channel subunit, alpha2/delta2), a methamphetamine-induced hnRNP H target exhibiting changes in 3’UTR splicing and transcript levels following methamphetamine exposure. Pharmacological inhibition of CACNA2D2 with FDA-approved drug pregabalin (a.k.a. Lyrica®) reduced methamphetamine-induced locomotion in wild-type mice, suggesting CACNA2D2 is a hnRHP H target that directly influences methamphetamine locomotion.

## 2. MATERIALS AND METHODS

### 2.1 Mice

Wild-type mice were generated by breeding male mice heterozygous for a 16 bp deletion in Hnrnph1 leading to a frameshift mutation and premature stop codon on an isogenic C57BL/6J background to C57BL/6J wild-type females ordered from The Jackson Laboratory (#000664) as previously described (Yazdani et al., 2015). We had initially included Hnrnph1 heterozygous mutants in all components of this study; however, upon re-sequencing of the TALENs-targeted region, it was discovered that although the mutants retained a second alternate cut site for the restriction enzyme BstNI (not present in wild-type mice) and thus appeared heterozygous for the original 16 bp mutation on a gel, the mutation underwent partial DNA repair over the course of repeated backcrossing to maintain the line, being reduced to 3 bp. This alternate, partially repaired allele nearly replaced the original 16 bp mutation due to a population bottleneck in the colony. There was also a separate segregating mutation where an additional 1 bp had been deleted (17 bp total in size). Therefore, we removed the mutants from all aspects of the results and focused solely on the results from isogenic C57BL/6J wild-type littermates. Both female and male offspring (ranging from 50 – 100 days old at the start of the experiment) from this breeding scheme were used in the study. Mice were housed in same-sex groups of 2-5 in standard mouse cages in ventilated racks under standard housing conditions on a 12 h:12 h light:dark schedule with food and water supplied ad libitum. All protocols involving mice were written in accordance with the Guideline for the Care and Use of Laboratory Animals and were approved by Boston University’s IACUC committee.

### 2.2 Methamphetamine-induced locomotor activity followed by dissection of whole striatum and cross-linking immunoprecipitation (CLIP)

As described (Yazdani et al., 2015), on Days 1 and 2, all mice received a saline injection (10 ml/kg, i.p.) and were recorded for locomotor activity in Plexiglas chambers (40 cm length x 20 cm width x 45 cm height) for 1 h. On Day 3, mice received either saline (i.p.) again or methamphetamine HCl (2 mg/kg, i.p.; Sigma Aldrich, St. Louis, MO; Cat#1399001) and were recorded for locomotor activity for 30 min and the whole striata (left and right sides) were dissected from each mouse at 30 min post-injection (Ruan et al., 2020a). Dissected whole striata were stored in RNAlater (ThermoFisher Scientific, Waltham, MA; Cat#AM7020) following manufacturer’s instructions to stabilize the RNA and protein, followed by CLIP.

Striata from four mice were pooled per replicate (3 replicates per Genotype per Treatment) to provide a sufficient amount of RNA for CLIP. Each replicate used for CLIP-seq and RNA-seq was generated by pooling striata from 4 mice across multiple litters. Each pool consisted of samples from 2 females and 2 males. The striatum was chosen because of its involvement in the methamphetamine locomotor stimulant response, reinforcement and reward (Keleta and Martinez, 2012; Lominac et al., 2014). Given the large amount of striatal tissue needed for generating the CLIP-seq libraries in this study, we did not examine the ventral tegmental area (VTA, where the nuclei of the dopaminergic neurons reside) because the number of mice required to obtain a sufficient amount of tissue from the VTA per replicate was not feasible. The striatum punches contained primarily post-synaptic cells, but they also contained presynaptic dopaminergic terminals originating from VTA as well as other presynaptic terminals (e.g., glutamatergic terminals from prefrontal cortex). Thus, any low levels of cytosolic hnRNP H (Wall et al., 2020) and its bound RNAs at the presynaptic terminals were expected to be captured within our samples and thus, potentially relevant to rapid, dynamic regulation of in the cellular response to methamphetamine. Tissue was flash frozen in mortar-filled liquid nitrogen and crushed into powder with the pestle and kept on dry ice in a 100 mm Petri dish until use. Prior to crosslinking, a portion of the pooled tissue from each replicate was removed and stored in −80°C for later RNA extraction and bulk RNA-seq library preparation. The tissue was kept on dry ice in the dish while crosslinking was performed for three rounds using a 400 mJ/cm^2^ dosage of 254 nm ultraviolet radiation. The crosslinked tissue was then homogenized in 1 ml of lysis buffer (50 mM Tris-HCl pH7.4, 100 mM NaCl, 1% NP-40, 0.1% SDS, 0.5% sodium deoxycholate (protect from light), protease and phosphatase inhibitor cocktail (1:100; ThermoFisher Scientific, Waltham, MA; Cat#78444) and recombinant RNasin ribonuclease inhibitor (1 ul/ml; Promega, Madison, WI; Cat#N2511) with a mechanical homogenizer followed by addition of TURBO DNase (2 μl; ThermoFisher Scientific, Waltham, MA; Cat#AM2239). The lysate was kept on ice for 30 min followed by the addition of RNAse I_f_ (125 U/ml; NEB, Ipswich, MA; Cat#M0243L) and was allowed to incubate in a thermomixer set to 1200 RPM at 37^°^C for 3 min. The lysate was then centrifuged at 20,000 x g for 20 min at 4°C and kept on ice until use.

For RNA-immunoprecipitation, 133.3 μl of MyOne Streptavidin T1 beads (Invitrogen, Waltham, MA; Cat#65602) were incubated with 5.8 μl of 1 mg/ml of Pierce^TM^ biotinylated Protein G (ThermoFisher Scientific, Waltham, MA; Cat#29988) and 20 μg of either the hnRNP H antibody (Bethyl, Montgomery, TX; Cat#A300-511A) or the rabbit IgG antibody (EMD Millipore, Burlington, MA; Cat#12-370) for 1 h. The antibody-coupled beads were washed five times with 0.5% IgG-free BSA (Jackson ImmunoResearch, West Grove, PA; Cat# 001-000-162) in 1X PBS, followed by three washes in lysis buffer. The lysate that was clarified by centrifugation was added to the coated and washed beads and incubated with end-to-end rotation at 4°C for 2 h followed by 2X wash with 500 μl of wash buffer (end-over-end rotation at 4°C for 5 min each) and another 2X wash with 500 μl of high salt wash buffer (end-over-end rotation at 4°C for 5 min each). The components of the wash buffer are as follows: 20 mM Tris-HCl pH7.4, 10 mM MgCl_2_, 0.2% Tween-20, and recombinant RNasin ribonuclease Inhibitor (1 uL/ml, Promega, Cat#N2511). The components of the high salt wash buffer are as follows: 50 mM Tris-HCl pH7.4, 1M NaCl, 1 mM EDTA, 1% NP-40, 0.1% SDS, 0.5% sodium deocxycholate, and recombinant RNasin ribonuclease inhibitor (1 uL/ml; Promega, Madison, WI; Cat#N2511). These washes were followed by an additional wash with 500 μl each of both wash buffer and high salt wash buffer. Two additional final washes were performed with 500 μl of wash buffer. The rest of the wash buffer was removed, and beads resuspended in 20 μl of 1X Bolt LDS non-reducing sample buffer (ThermoFisher Scientific, Waltham, MA; Cat#B0007). The beads in sample buffer were then heated at 70°C for 10 min prior to SDS-PAGE on a 4-12% gradient NuPAGE Bis/Tris gels (ThermoFisher Scientific, Waltham, MA; Cat#NP0322BOX) followed by transfer to nitrocellulose membrane with 10% methanol for 6 hours at constant 150 mA. The radiolabeling experiments for determining the optimal CLIP conditions were performed as described in Rieger et al., 2020.

### 2.3 CLIP-seq and total RNA-seq library preparation

Following the completion of membrane transfer, the membrane was cut with a clean razor to obtain a vertical membrane slice per lane from 50 kDa (molecular weight of hnRNP H) to 75 kDa, which translates to 30 to 70 nucleotide RNA fragments crosslinked to the protein. RNA was then extracted from the membrane slices following a previously described procedure (Rieger et al., 2020). The same extraction procedure (starting with the addition of 7M urea) was used to isolate RNA from samples previously stored for total RNA-seq. Following RNA extraction, sample concentration was quantified with Agilent Bioanalyzer and approximately 0.2 ng of RNA was used to prepare next-generation sequencing libraries following the eCLIP procedure (Van Nostrand et al., 2016) with minor modifications according to Rieger et al., 2020. Because the RNA adapter on each sample contained a unique barcode, the cDNA libraries generated from the sample (12 CLIP-seq and 12 RNA-seq libraries) were multiplexed and pooled for increased throughput. A pooled library at a concentration of 10 nM was shipped to the University of Chicago Sequencing core and subjected to 100 bp paired-end 2 x 100 sequencing in a single lane on Illumina HiSEQ4000. To increase read coverage for the RNA-seq samples, those 12 cDNA libraries were pooled (10 nM concentration) for sequencing for a second time in one lane using Illumina HiSEQ4000.

### 2.4 CLIP-seq and total RNA-seq data processing

Reads were trimmed for quality using Trimmomatic (Bolger et al., 2014). Unique Molecular Identifier (UMI) sequences were extracted from Read 2 for removal of PCR amplification duplicates using the ‘extract’ command in UMI-tools (Smith et al., 2017). After the reads were trimmed and UMI extracted, we used STAR (Dobin et al., 2013) to map reads to the mouse genome, version GRCm38/mm10. Even though the samples had undergone ribosomal RNA (rRNA) depletion prior to library preparation, we used BEDtools (Quinlan and Hall, 2010) to intersect the bam files with the RNA annotation bed file exported from UCSC Table browser (Karolchik et al., 2004) to remove rRNAs and other repetitive RNAs. Using the ‘dedup’ command in UMI_tools (Smith et al., 2017), PCR duplicates from the rRNA-depleted bam files were removed based on UMI extracted in the previous step.

### 2.5 Peak calling using <u>CL</u>IP-seq <u>A</u>nalysis of <u>M</u>ulti-mapped reads (CLAM)

To define the binding sites for hnRNP H for each of the four conditions separately, we used deduplicated bam files for the three replicates of each condition for peak calling in CLAM (Zhang and Xing, 2017). The BAM file for the CLIP sample and the BAM file for input (corresponding RNA-seq sample) were used as input along with a gene annotation file in Gene Transfer Format downloaded from GENCODE. Following the steps described in CLAM (Zhang and Xing, 2017), in order to allow for peak calling of multi-mapped reads, BAM files were preprocessed (‘preprocessor’ command) first to separate multi-mapped reads and uniquely mapped reads followed by realigning (‘realigner’ command) and then peak calling (‘peakcaller’ command) on a gene-by-gene basis in 100-bp bins. The called peaks were then annotated to the genomic regions using the ‘peak_annotator’ command to examine binding site distributions across the 5’UTR, 3’UTR, CDS, and introns.

### 2.6 Homer *de novo* Motif Discovery

To identify the top over-represented motifs in the peaks (those peaks with CLIP peak intensity > 1.5) that were identified in CLAM, Homer software (Heinz et al., 2010) was used for *de novo* motif discovery by using “findMotifsGenome.pl” function. The input file comprised a list of hnRNP H associated peaks containing the genomic coordinates. The “annotatePeak.pl” function was then used to identify motif locations to find genes containing a particular motif.

### 2.7 Gene Ontology and KEGG Pathway Enrichment Analysis

To examine the biological function of the hnRNP H targets (peak signal value defined by a CLAM value greater than 1.5 for each target), we performed gene ontology (GO) and KEGG pathway enrichment analysis of the significant targets using GOseq (Young et al., 2010), which normalizes bias toward long transcripts. The GO terms and KEGG pathways that met the threshold (adjusted p values < 0.05) were organized into networks for clear visualization and easy interpretation in Cytoscape using the ClueGo plug-in (Bindea et al., 2009).

### 2.8 Alternative Exon Usage Analysis using real-time quantitative PCR (RT-qPCR)

Striatal tissue for RT-qPCR validation was collected 30 min after saline or methamphetamine injections as described above. The left striatum was dissected as previously described (Ruan et al., 2020a) and subsequently stored in RNAlater (ThermoFisher Scientific, Cat#AM7020) following manufacturer’s instructions to stabilize the RNA. Following RNA extraction with TRIzol reagent (Invitrogen, Cat#15596026), oligo-DT primers were used to synthesize cDNA from the extracted total RNA using a high-capacity cDNA reverse transcription kit (ThermoFisher Scientific, Cat#4368814). RT-qPCR using PowerUP SYBR Green (ThermoFisher Scientific, Cat#A25741) was then performed to evaluate alternative 3’UTR exon usage of Cacna2d2 or overall differential expression of Cacna2d1 (last exon junction near the 3’ end of the gene spanning all UCSB-annotated transcripts). The primer sequences are listed here:

– Proximal end of the 3’UTR of *Cacna2d2* (forward primer): 5’-TTGGCCACTCTCTCCTGAAG-3’

– Proximal end of the 3’UTR of *Cacna2d2* (reverse primer): 5’-ACTAGTGGCCTCCTGTCCTA-3’

– Distal end of the 3’UTR of *Cacna2d2* (forward primer): 5’-CCCCATCAGGTAGTTGTCCA-3’

– Distal end of the 3’UTR of *Cacna2d2* (reverse primer): 5’-TGTCGCTGTTGTTTTCCCAA-3’

– Exons 38-39/proximal 3’UTR of *Cacna2d1* (forward primer): 5’-GCAGCCCAGATACCGAAAAG-3’

– Exons 38-39/proximal 3’UTR of *Cacna2d1* (reverse primer): 5’-TCGTTGCAGATCTGGGTTCT −3’

– Exons 5-6 of *Gapdh* (forward primer): 5’-GCCTTCCGTGTTCCTACC-3’

– Exons 5-6 of *Gapdh* (reverse primer): 5’-CCTCAGTGTAGCCCAAGATG-3’

### 2.9 Effect of Pregabalin Pre-treatment on Locomotor Responses following Saline and Methamphetamine Treatment and Striatum Dissections

To habituate mice to the testing environment, on Days 1 and 2, mice were pretreated with a saline injection (10 ml/kg, i.p.) and placed back into their home cages. 30 min later, mice were administered a second saline injection (10 ml/kg, i.p.) and were recorded for locomotor activity in Plexiglas chambers (40 cm length x 20 cm width x 45 cm height) for 1 h. On Day 3, mice were pretreated with either saline (10 ml/kg, i.p.) or pregabalin (30 mg/kg, i.p., Biosynth Carbosynth, San Diego, CA; Cat#FA27139). This pregabalin dose was previously shown to attenuate cocaine intravenous self-administration (de Guglielmo et al., 2013) without inducing addiction model behaviors on its own (Coutens et al., 2019). All mice within the same cage received the same pretreatment. 30 min later, mice were injected with methamphetamine (2 mg/kg, i.p.). 60 min following methamphetamine (or saline) and 90 min post-pregabalin injections, whole striata (left and right sides) were dissected from each mouse (Ruan et al., 2020a). Because pregabalin inhibits CACNA2D2 and because Cacna2d2 is an hnRNP H target, we wanted to know if pregabalin pre-treatment would lead to a cellular adaptive response as measured via changes in Cacna2d2 and Cacna2d1 transcript levels following subsequent saline versus methamphetamine treatment. Dissected left striata were stored in RNAlater following manufacturer’s instructions to stabilize the RNA and for quantification of *Cacna2d2* 3’UTR usage and Cacna2d1 expression.

### 2.10 Experimental Design and Statistical Analyses

#### Locomotor activity in 5-min bins

To determine the effect of pregabalin pretreatment on methamphetamine-induced locomotor activity, total distance traveled in 5-min bins was analyzed separately for Days 1, 2, and 3 using mixed effects ANOVA with pre-Treatment (saline, pregabalin) as the between-subject factor and Time as the repeated-measure factor. Main Treatment (saline, methamphetamine) was not included as these mice were run in separate cohorts. Therefore, all stats were conducted within each main Treatment group. For Day 3, the significant two-way (Pre-Treatment x Time) interaction was followed up with Bonferroni-adjusted multiple pairwise comparisons to identify difference between saline versus pregabalin in each 5-min time bin.

#### Effect of Methamphetamine on hnRNP H binding targets

Deduplicated BAM files were merged across all CLIP samples and deduplicated BAM files were merged across all input (bulk RNA-seq) samples to generate two merged BAM files as the input for peak calling in CLAM as described above. The called peaks were then annotated to the genomic regions as described above. The file containing all of the peaks was formatted into a file in GTF format that was used for strand-specific feature counting in individual BAM files from the CLIP samples using featureCounts in SubRead (Liao et al., 2014). Only peaks with peak intensity > 1.5 and FDR < 0.05 as defined by CLAM were included as ‘features’ for read counting. Reads were summed for each significant peak to produce a count table containing peaks as rows and samples in columns for differential analysis. To examine the effect of methamphetamine on hnRNP H binding, differential analysis was performed using DESeq2 (Love et al., 2014) based on summed read counts for each peak derived from Subread featureCount. Differential hnRNP H binding to *CacnA2D2* specifically was also performed with a t-test on normalized counts.

#### Differential gene expression and differential exon and intron usage analyses

To triangulate on the CLIP-seq and RNA-seq datasets and associate the downstream effects of hnRNP H binding on gene expression and alternative splicing, we analyzed differential gene expression using DESeq2 and differential exon/intron usage using ASpli (Mancini et al., 2020). Deduplicated BAM files from the total RNA-seq samples were used as inputs for overall gene expression analysis and alternative splicing. Comparing differential expression and exon/intron 3’UTR usage for *Cacna2d2* specifically was also performed using t-tests on the normalized counts.

#### Validation of differential exon usage of the 3’UTR of Cacna2d2

For analyzing differential 3’UTR proximal versus distal usage of *Cacna2d2* or overall differential expression of *Cacna2d1*, the fold-change for each condition relative to saline-treated wild-types was calculated using the 2^-ΔΔCT^ method (Livak and Schmittgen, 2001). Samples were plated in triplicate as technical replicates. Significance of the RT-qPCR results was determined using t-tests.

### 2.11 Code/Software

Codes associated with analyses mentioned in Method Details can be found at: https://github.com/camronbryant/hnrnph1_clip.

## 3. RESULTS

### 3.1 hnRNP H binding sites contain G-rich binding motifs that are enriched in the intronic regions

Accumulating evidence from our lab indicates a role for hnRNP H in methamphetamine addiction liability (Ruan et al., 2020a; Borrelli et al., 2021). Defining the set of striatal RNA targets in brain tissue that are differentially regulated by hnRNP H at baseline versus in response to methamphetamine could provide new insight into the cellular and molecular mechanisms underlying methamphetamine-induced dopamine release and behavior. To identify the *in vivo* hnRNP H targets, we performed CLIP using an antibody specific for hnRNP H in the striatum harvested 30 min following saline or methamphetamine treatment. The experimental design as described in Section 2.2, along with the read coverage per pooled sample is outlined in **Table S1**. We previously validated this antibody to be specific for the C-terminus of hnRNP H via immunoadsorption with a blocking peptide for the epitope (Ruan et al., 2018). Here, we show that the antibody specifically pulled down hnRNP H at 50 kDa with no signal detected using rabbit IgG (**Figure S1A**; Lane 6 versus Lane 3). Importantly, we used stringent lysis and wash conditions as published (Van Nostrand et al., 2016). Twenty μg of hnRNP H antibody was the optimal amount needed as indicated by visual inspection of the band intensity (**Figure S1B**).

Our CLIP procedure resulted in RNA-specific pulldown of hnRNP H-RNA complexes. RNAse fragmented the CLIP samples into different sizes in a concentration-dependent manner (**Figure 1A**; from left to right, lanes 3 – 6) indicating the hnRNP H CLIP was RNA-dependent. Longer time periods of incubation of CLIP samples with RNase yielded a lower amount of RNA, providing further support that hnRNP H CLIP was RNA-dependent (**Figure S1C**; Lanes 2-5). Negative controls included immunoprecipitation (**IP**) from uncrosslinked sample (**Figure 1A**; Lane 1) and immunoprecipitation using rabbit IgG from wild-type striatal tissues (**Figure 1A**; Lane 2). No RNA was detected in these two negative control samples (**Figure 1A**; lanes 1 and 2), indicating both the need for UV-crosslinking for RNA pull down and the specificity of hnRNP H pull down. We chose a region 30 – 70 nucleotides in size (50 – 80 kDa) for RNA extraction to capture the targets of hnRNP H *in vivo* (**Figure 1A**). The cDNA libraries generated from the CLIP samples of the IgG IPs did not yield any detectable PCR bands using gel electrophoresis (only dimers from the sequencing adapter libraries), even after 28 PCR cycles (**Figure S2A**). For this reason, none of the four IgG cDNA libraries were subjected to sequencing. However, for the CLIP samples, DNA bands corresponded to the correct size of the cDNA library (>150 bp) and were detected after 18 PCR cycles (**Figure S2B**). We subjected the same samples used in CLIP for total RNA-seq and measured starting transcript abundance, which permitted normalization to account for differences in RNA abundance indicated by read counts (**Table S1**).

**Figure 1.**
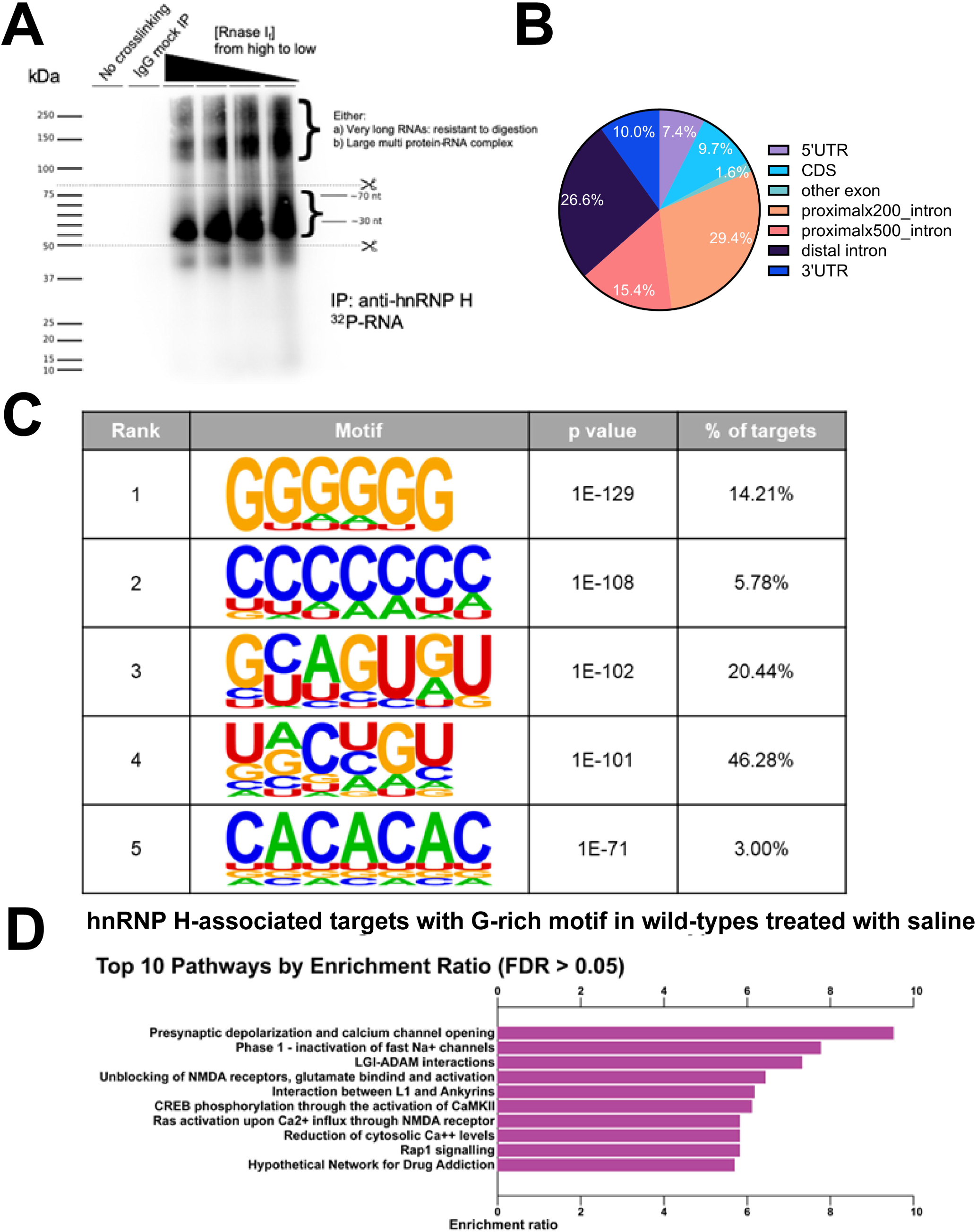
Striatal RNA-binding sites of hnRNP H at baseline (saline control) are enriched for introns and G-rich binding motifs. eCLIP-seq revealed transcriptome-wide striatal RNA targets of hnRNP H in saline control wild-types (WT_SAL). Peak calling of both uniquely and multi-mapped reads was performed using CLAM (Zhang and Xing, 2017). **(A):** RNA ^32^P autoradiogram of hnRNP H-bound RNA. CLIP conditions are shown for each lane: no crosslinking, IgG mock IP, and four different RNase I_f_ concentrations from high to low. As expected, an increasing amount of RNA was pulled down as the concentration of RNase I_f_ was decreased. The scissors denote the region above the molecular weight of hnRNP H (50 kDa) that was isolated for sequencing. The region runs from 50 to 80 kDa, which corresponds to 30 to 70 bp for RNA length. The larger bands that were greater than 100 kDa could represent very long RNAs that are resistant to digestion or large hnRNP H-associated protein-protein complexes bound to RNAs. **(B):** hnRNP H binds primarily to introns of target RNAs. More than 70% of hnRNP H CLIP sites are intronic. The pie chart shows the relative distribution of CLIP sites in 5’UTR, coding sequences (CDS), introns, and 3’UTR. Proximal introns indicate less than 200 (proximalx200_intron) or 500 (proximalx500_intron) nucleotides from the 5’ or 3’ splice sites with the remainder annotated as distal introns. Unannotated exons are referred to as “other exons”. **(C):** Poly-G tract was the most prevalent component of the hnRNP H binding motif. *De novo* motif discovery of hnRNP H CLIP sites was performed using Homer (Heinz et al., 2010). **(D):** hnRNP H RNA-binding targets containing G-rich motifs in their binding sites were most highly enriched for “presynaptic depolarization and calcium channel opening” pathway. Pathway enrichment analysis of hnRNP H RNA-binding targets with the poly-G motif was performed in Goseq (Love et al., 2014) to account for transcript length bias. The top 10 pathways with FDR < 0.05 are shown, sorted from high to low enrichment ratio.

To define the targets of hnRNP H under basal conditions, we first focused our analysis on baseline saline treatment. Specific hnRNP H sites were identified using Peakcaller subcommand in CLAM (Zhang and Xing, 2017) to perform peak calling throughout the whole genome. The peak calling process was done on a gene-by-gene basis by breaking down each gene into 100 nucleotide bins and testing for enrichment of mapped reads over control (or total RNA-seq mapped reads) and specifying a negative-binomial model on observed read counts. Importantly, CLAM calls peaks using a combination of uniquely- and multi-mapped reads for inclusion of RNA-binding sites that are repetitive elements (Zhang and Xing, 2017). hnRNP H-associated peaks were defined as enriched (p < 0.05 and peak signal intensity >1.5) in CLIP samples over input RNA-seq samples. The genomic region annotated peaks for each saline and methamphetamine can be found in the **Table S2** and **Table S3**, respective for treatment. Analysis of the peaks across gene subregions revealed enriched intronic binding of hnRNP H, comprising about 70% of total distribution (**Figure 1B**). This finding is consistent with previous characterization of hnRNP H in HeLa cells (Huelga et al., 2012; Uren et al., 2016), supporting successful isolation of hnRNP H-bound RNAs in mouse striatal tissue.

*De novo* motif discovery of significant hnRNP H-associated binding sites using the Homer database (Heinz et al., 2010) detected the top over-represented motif to be G-rich (**Figure 1C**) and was more prevalent in intronic regions and 3’ UTRs (**Table S4**) which agrees with the prior literature (Lefave et al., 2011; Uren et al., 2016) and indicates that our CLIP procedure successfully isolated hnRNP H targets in mouse brain striatal tissue. Intronic poly G-stretches can enhance hnRNP H binding (Han et al., 2005) and their length predicts splicing (Katz et al., 2010). Pathway enrichment analysis of the hnRNP H RNA targets containing these G-rich genomic locations identified “presynaptic depolarization and calcium channel opening,” comprising of subunits of calcium channels encoded by *Cacna1a, Cacnb4, Cacna1e, Cacng4, Cacna1b, Cacng2, and Cacnb2* (**Figure 1D and Table S4**), all of which are important for neurotransmission (Dolphin and Lee, 2020). FMRP is an RBP that was previously shown to bind to mRNA transcripts that encode for ion channels (Darnell et al., 2011). Thus, like FMRP, our findings identify hnRNP H as another RBP that binds to calcium channel subunit transcripts to regulate gene splicing and expression that plausibly influences neurotransmission and behavior (**Tables S5 and S6**).

### 3.2 Methamphetamine treatment induces changes in RNA-binding targets of hnRNP H

To test the hypothesis that hnRNP H exhibits a functional molecular response to methamphetamine treatment (i.e., a change in the hnRNP H targetome), we examined the baseline and methamphetamine-induced targetome of hnRNP H. Comparing changes in binding across subregions of the mRNA transcripts revealed the extent to which methamphetamine could impact splicing versus mRNA stability and/or translation. The positions of the peaks and the corresponding gene targets for effect of methamphetamine are provided in **Table S7.**

Analyzing the read distribution separately across the 3’UTR, introns, CDS, and 5’UTR for methamphetamine versus saline, we again discovered over-representation of binding sites for intronic regions and 3’ UTRs (see below) of the mRNA transcripts across the two conditions (**Figure 2A**). Thus, methamphetamine treatment induced a signaling response that ultimately shifted the binding of hnRNP H to its intronic targets that are predicted to perturb mRNA splicing. Notably, following methamphetamine, there is an *increase* in the percentage of 3’UTR binding sites (**Figure 2A and Table S8**). These results suggest that methamphetamine-induced signaling within the striatum induces alterations in hnRNP H-mediated regulation of mRNA stability, polyadenylation site usage, and/or translation.

**Figure 2.**
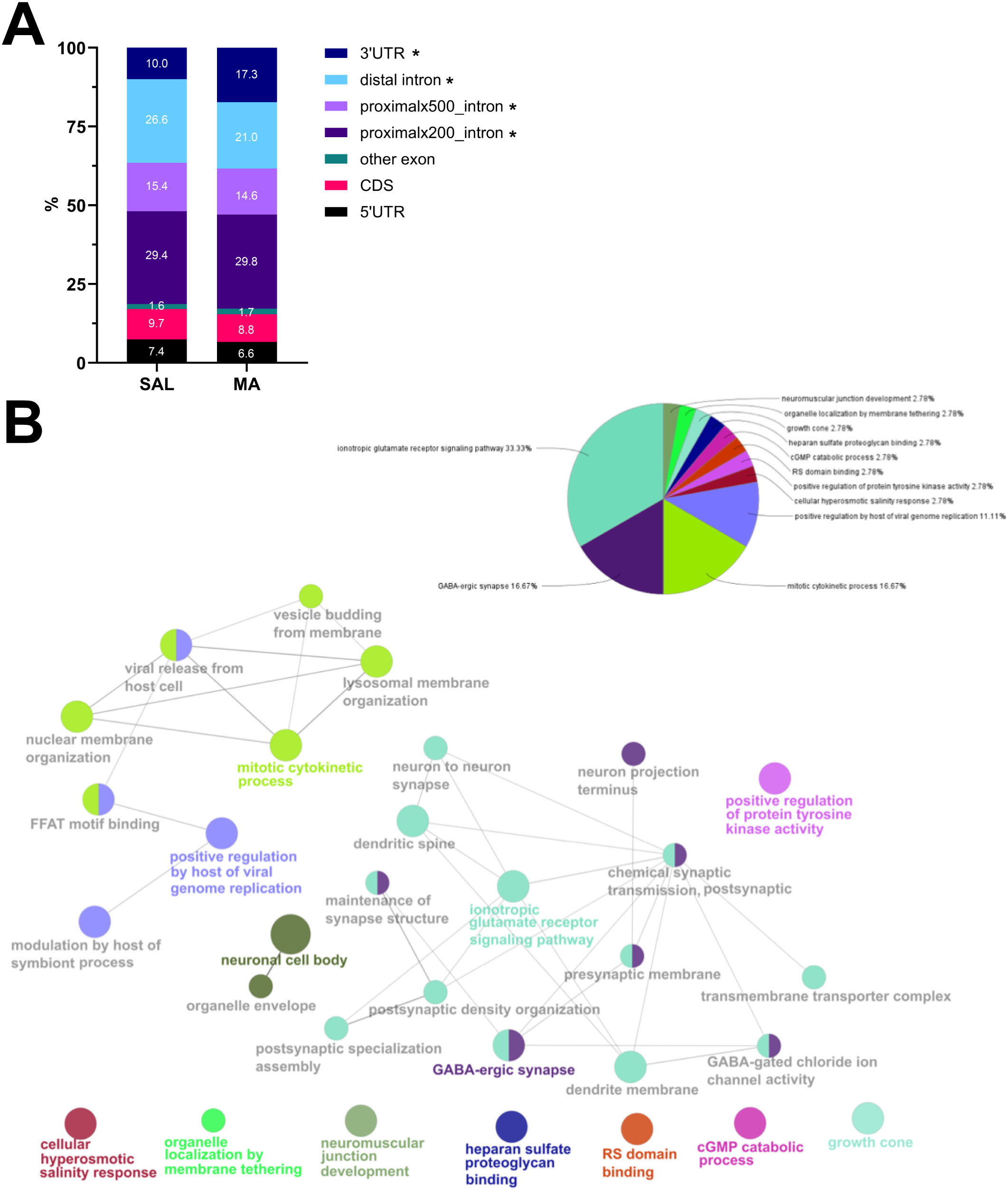
Methamphetamine-induced striatal RNA-binding targets of hnRNP H are enriched for pathways and cellular components involved in drug-evoked synaptic plasticity. CLIP-seq analysis revealed transcriptome-wide striatal RNA-binding targets of hnRNP H at 30 min post-saline (i.p.) versus methamphetamine treatment (2 mg/kg, i.p.). Peak calling in CLAM (Zhang and Xing, 2017) was performed separately for both conditions. For differential analysis, peak calling was also performed using CLAM (Zhang and Xing, 2017) on the merged bam file across the two conditions followed by read counts against the identified peaks and differential analysis of peak intensity. SAL = saline; MA = methamphetamine. **(A):** In comparison to SAL, only the percentages of hnRNP H binding events associated with 3’UTR and introns were significantly different in the other three conditions (See chi-square tests in **Table S5**). The percentage of binding events that comprised 3’UTRs varied robustly between SAL vs MA (chi-square test: *p < 0.001). All intron binding events (including distal introns and proximal introns) were significantly different from SAL (chi-square test: all p’s < 0.001), The relative distribution of RNA-binding sites over the gene elements for each condition is shown. **(B):** GO enrichment and pathway enrichment analysis using Goseq were conducted on differential hnRNP H CLIP targets between MA vs. SAL. The significant GO and KEGG terms showing FDR-adjusted p-value of less than 0.05 are annotated by ClueGO to create a functionally organized GO/pathway term network shown. The Venn diagram summarizes the different functional terms in the network with “ionotropic glutamate receptor signaling pathway”, “GABA-ergic synapse”, and “mitotic cytokinetic process” being the top 3 leading terms.

Our CLIP-seq dataset identified more than 1000 mRNA transcripts associated with methamphetamine effects. The enrichment map network of the top over-represented pathways and GO terms for these targets revealed clustering of nodes associated with synaptic function (**Figure 2B and Table S9**). Post-transcriptional regulation of these mRNA targets by hnRNP H could contribute to reduced methamphetamine-induced dopamine release and/or reduced post-synaptic dopaminergic signaling.

### 3.3 3’UTR targets of hnRNP H show enrichment for excitatory and psychostimulant-induced synaptic plasticity

Methamphetamine treatment modulated *Hnrnph1* binding events within 3’UTRs (**Figure 2A**). Given the proximity between 3’UTRs and protein translation and stability, we reasoned that those methamphetamine-mediated 3’ UTR targets in hnRNP H binding are potential mechanistic targets of the cell biological response altering synaptic transmission and behavior and thus warranted further investigation. We partitioned the hnRNP H-interactive CLIP peaks into separate subgenic regions (5’UTR, 3’UTR, intron, or CDS), which allowed us to characterize the impact of changes in binding on specific types of hnRNP H-dependent post-transcriptional processing that could refine which subgenic targets are most relevant in driving the enrichment scores related to synaptic function. We examined binding of hnRNP H across transcripts by plotting normalized log_2_CPM (counts per million) mapped reads to subgenic regions (**Figure 3**). We observed an overall increase in hnRNP H binding to the 3’UTR region of the detected RNA targets in response to methamphetamine and this pattern was most pronounced for the most robust 3’UTR targets as illustrated by the heat map (**Figure 3**). Binding of RBPs to 3’UTRs of transcripts provides a rapid means for regulating mRNA translation at sites distant from the cell body such as transcripts for synaptic proteins (Mayr, 2017; Harvey et al., 2018). Thus, in response to methamphetamine, hnRNP H could rapidly regulate translation of proteins involved in synaptic function via changes in 3’UTR binding to mRNA transcripts.

**Figure 3.**
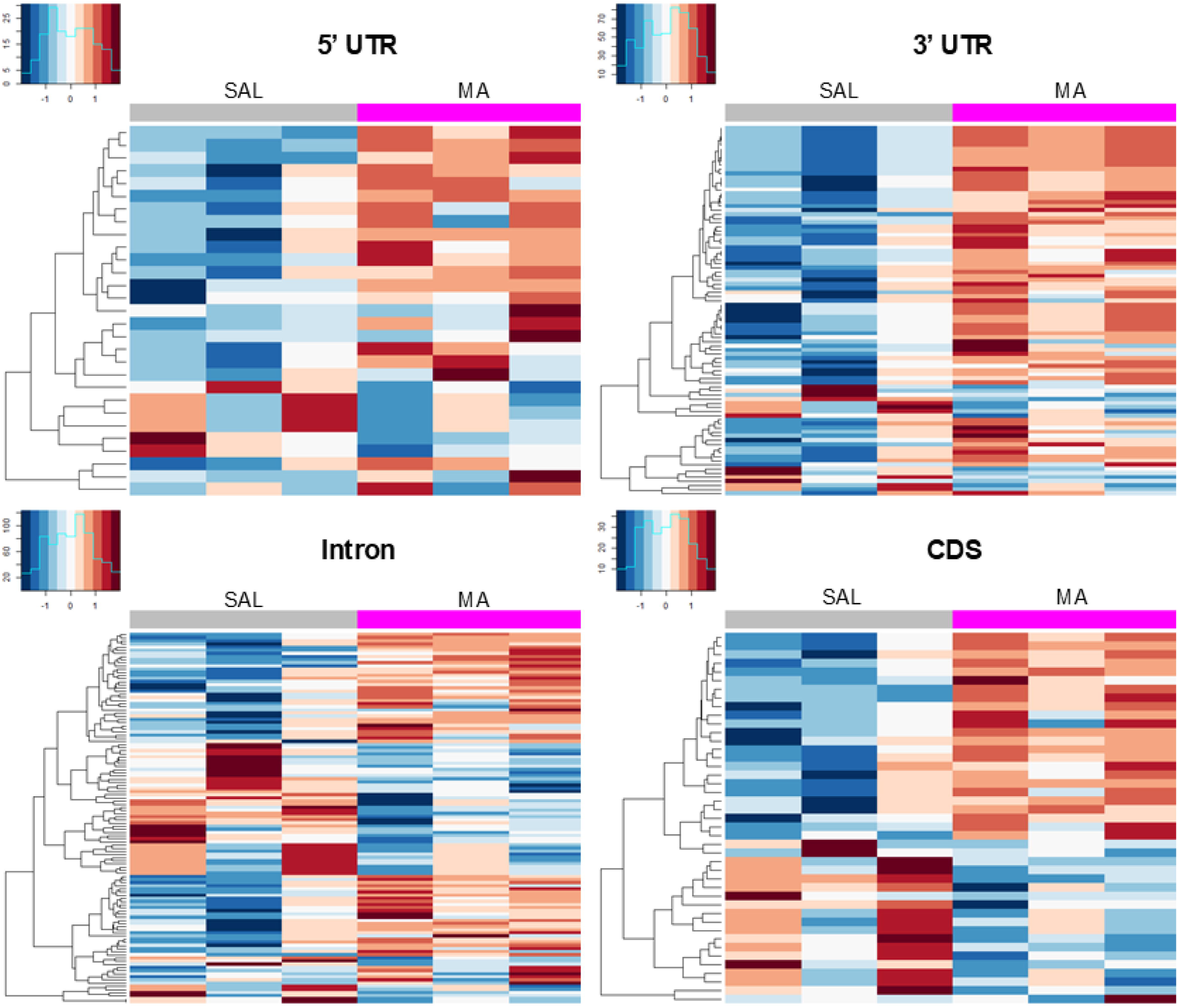
3’UTR targets of hnRNP H show a methamphetamine-induced increase in binding in striatum. Methamphetamine-induced hnRNP H CLIP peaks, stratified across four gene elements, including 5’UTR, 3’UTR, intron, and CDS. SAL = saline; MA = methamphetamine. Heatmaps show normalized log_2_ CPM (count-per-million) read counts for significant peaks showing a MA treatment effect. The row of each heatmap represents a binding site sorted by p-value from smallest to largest, with those with a p-value of less than 0.05 shown in the heatmap. Each column represents a sample, with the first three samples (grouped by the grey bar) being SAL-treated and the final 3 samples (grouped by the pink bar) being MA-treated. In response to MA, there was an increase in baseline binding of hnRNP H to the 3’UTR regions of RNA-binding targets.

### 3.4 Methamphetamine treatment induce changes in RNA-binding targets of hnRNP H

We next sought to determine changes in mRNA transcript levels associated with methamphetamine-induced changes in mRNA binding. To integrate striatal hnRNP H binding with gene expression and alternative splicing, we analyzed the transcriptome of striatal tissue from the same samples used in CLIP-seq. We performed both gene- and exon/intron-level transcriptome analyses to identify differentially expressed genes and genes showing evidence for alternative splicing, namely genes showing significant differential exon or intron usage. The outputs from DESeq2 differential gene expression analysis and ASpli differential splicing analysis are provided in **Table S10** and **Table S11**. There were 7 genes (*Agrn, Arhgef26, Dctn1, Kalrn, Litaf, Ptprs, Ubqln1)* that showed differential binding and alternative splicing, 2 genes (*Mid1, Elp6*) that showed differential binding and gene expression, and one gene *(Piezo1)* that showed differential expression and differential splicing in response to methamphetamine **(Fig 4A-B).**

**Figure 4.**
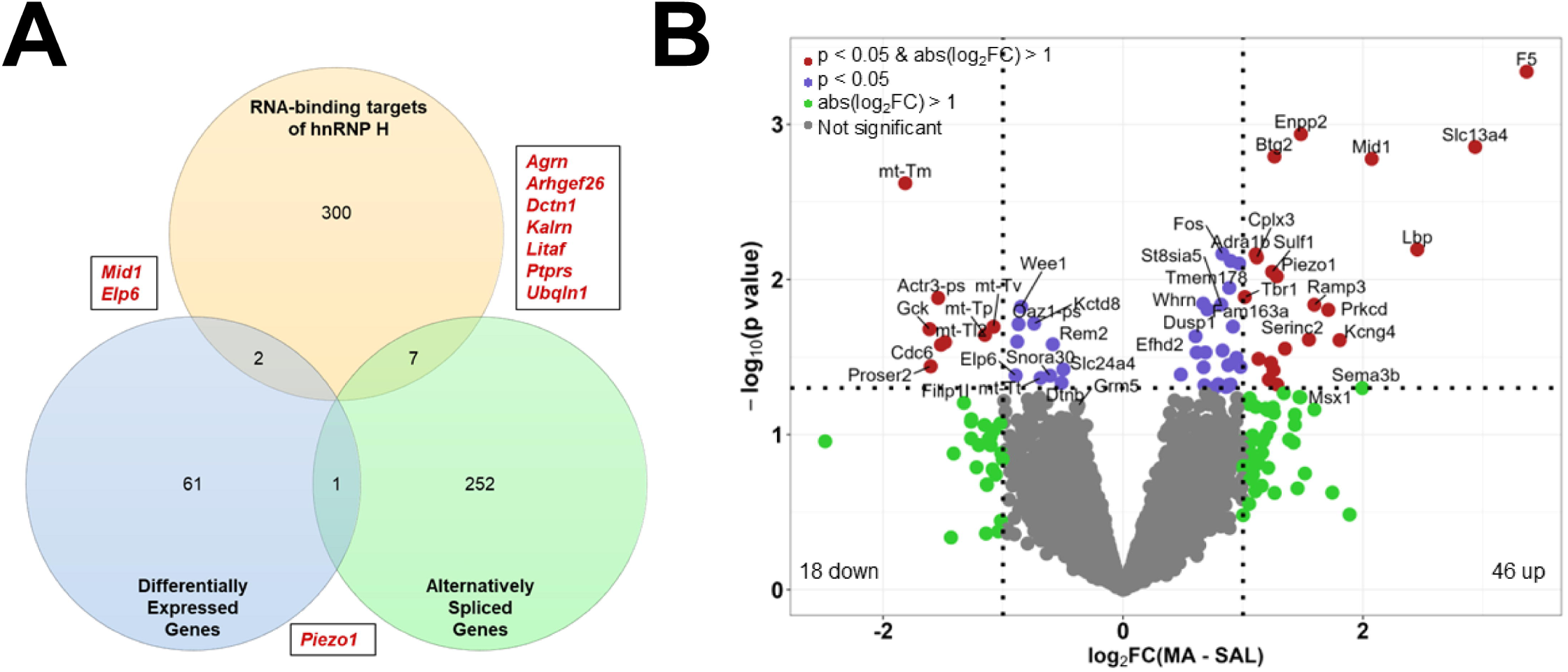
RNA targets showing a methamphetamine-induced change in hnRNP H1 binding, gene expression, and alternative splicing in striatum. Differential gene expression (DE) comparing MA vs SAL was performed using DESeq2 (Love et al., 2014). Differential exon and intron usage analysis (alternatively spliced; AS) for the effect of methamphetamine was performed using Aspli (Mancini et al., 2020). SAL = saline; MA = methamphetamine. **(A):** Venn diagram comparing RNA-binding targets of hnRNP H with differential gene expression and differential exon or intron usage. The 64 DE genes and 260 AS genes were compared with the 327 hnRNP H targets that showed differential binding in response to methamphetamine. The overlapping genes are highlighted in red. **(B):** Volcano plot of differentially expressed genes in response to methamphetamine.

Our prior targetome, transcriptome, and spliceome analysis that had included Hnrnph1 mutants versus wild-types triangulated on *Cacna2d2* as a candidate gene that showed Genotype x Treatment effects and could mediate methamphetamine-induced synaptic transmission and behavior (Ruan *et al*., 2021). CLIP analysis indicates that hnRNP H binds to the 3’UTR of *Cacna2d2* and thus could regulate synaptic localization, polyadenylation site selection, mRNA stability at the 3’UTR, and ultimately CACNA2D2 protein levels (**Figure 5A**). Three putative *Cacna2d2* isoforms, harboring different 3’UTR lengths, have been annotated by GENCODE. The MEME suite tools (Bailey et al., 2009) identified G-rich motifs (canonical for hnRNP H) within the 3’UTR of *Cacna2d2* (**Figure 5B**). Additionally, we observed a significant decrease in *Cacna2d2* gene expression in response to MA (p = 0.0295; **Figure 5C**) along with a trending decrease in H1 binding (p = 0.0889; **Figure 5D**) and visible increase in H1 binding for 2 of the 3 samples (p = 0.1419; **Figure 5E).** Together, these results warranted an in vivo behavioral pharmacological study to directly test the role of CACNA2D2 in methamphetamine-induced behavior.

**Figure 5.**
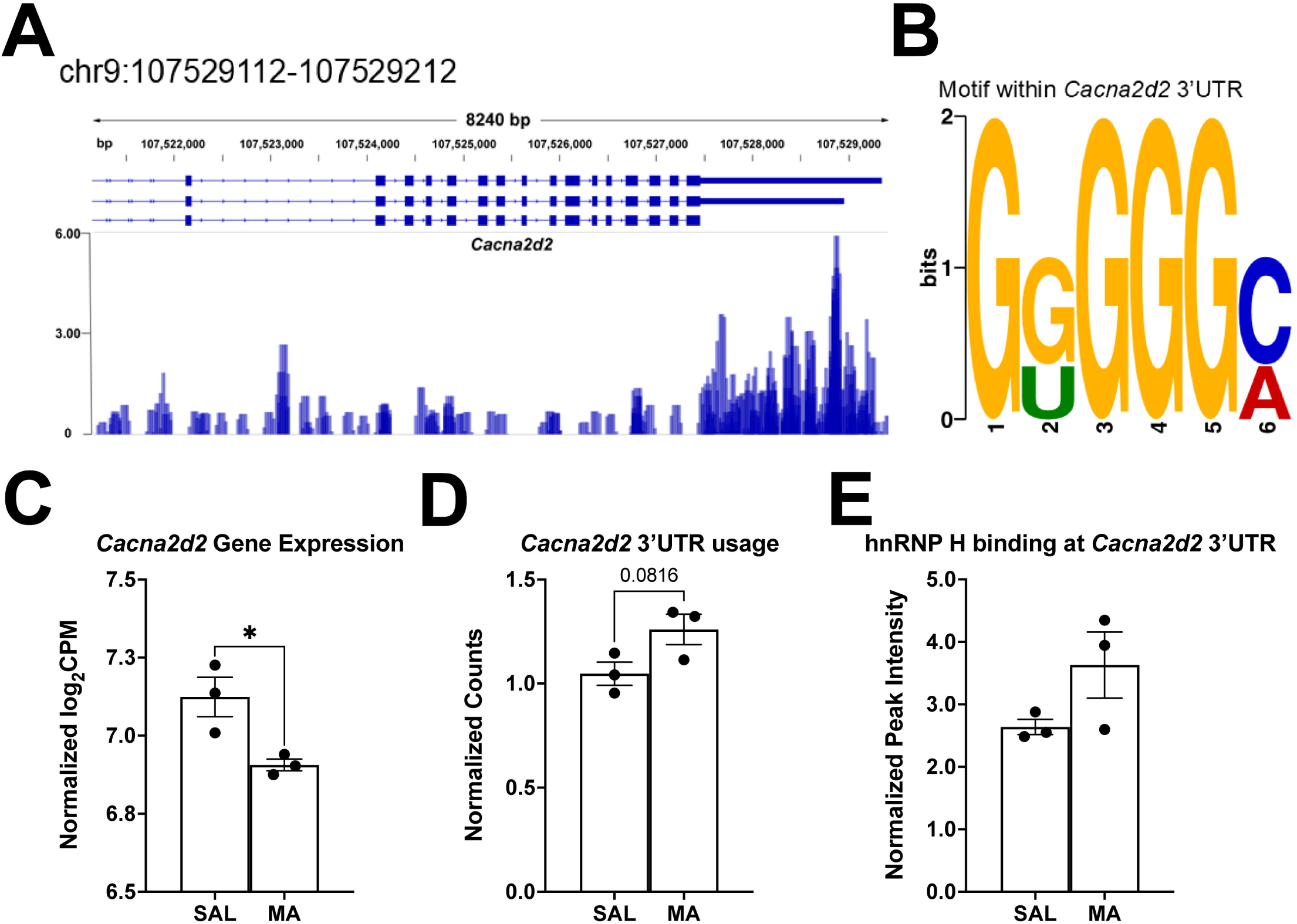
Methamphetamine-induced change in hnRNP H1 binding, gene expression, and alternative splicing of Cacana2d2 in striatum. **(A):** hnRNP H preferentially binds to the 3’UTR of *Cacna2d2*. Visualization of reads by Integrative Genome Browser (Thorvaldsdóttir et al., 2013) for the hnRNP H CLIP peak at the 3’UTR of *Cacna2d2*. Scale of the plot height is in counts per million (CPM). **(B):** A G-rich motif was detected at the hnRNP H CLIP peak within the 3’UTR of *Cacna2d2*. *De novo* motif discovery of the binding site was performed in MEME (Bailey et al., 2009). **(C):** Effect plots showing methamphetamine-induced decrease in hnRNP H binding to the 3’ UTR of *Cacna2d2* [left; t(4) = 2.346, p = 0.0789], increased usage of the 3’UTR [middle, t(4) = 2.315, p = 0.0816], and a statistically significant decrease in transcript levels of *Cacna2d2* (right, [t(4) = 3.317, p = 0.0295). Here, 3’UTR usage is defined as to the number of normalized reads mapped to the 3’UTR portion of *Cacna2d2*. The effect plots were generated from CLIP-seq and RNA-seq data showing average values with standard error of the means, with n = 3 per condition.

### 3.5 Pregabalin, a CACNA2D2/CACNA2D1 inhibitor

To test the contribution of CACNA2D2 to methamphetamine behavior, we selected pregabalin (PGB; a.k.a. Lyrica©), an FDA-approved drug that targets both CACNA2D2 and CACNA2D1. A timeline of pregabalin and methamphetamine injections, open field locomotor assessment, and tissue harvesting is presented in **Figure 6A**. Importantly, there was no spurious effect of random treatment assignment (PGB versus vehicle) on locomotion during saline habituation days in saline-treated animals **(Figure 6B-C).** Furthermore, on test day (Day 3), there was no effect of pregabalin on locomotion **(Figure 6D)**. In methamphetamine-treated animals, there was no spurious effect of random treatment assignment (PGB vs vehicle) on locomotion during saline habituation days **(Figure 6E-F)**. On test day (Day 3), pregabalin pretreatment induced a significant reduction in methamphetamine-induced hyperlocomotion (**Figure 6G**; p = 0.0430). There was also a significant Treatment x Time interaction (p = 0.0042), and while multiple comparisons with Bonferroni corrections did not identify any significant time windows, the greatest decrease was localized between 15-35 min into the session.

**Figure 6.**
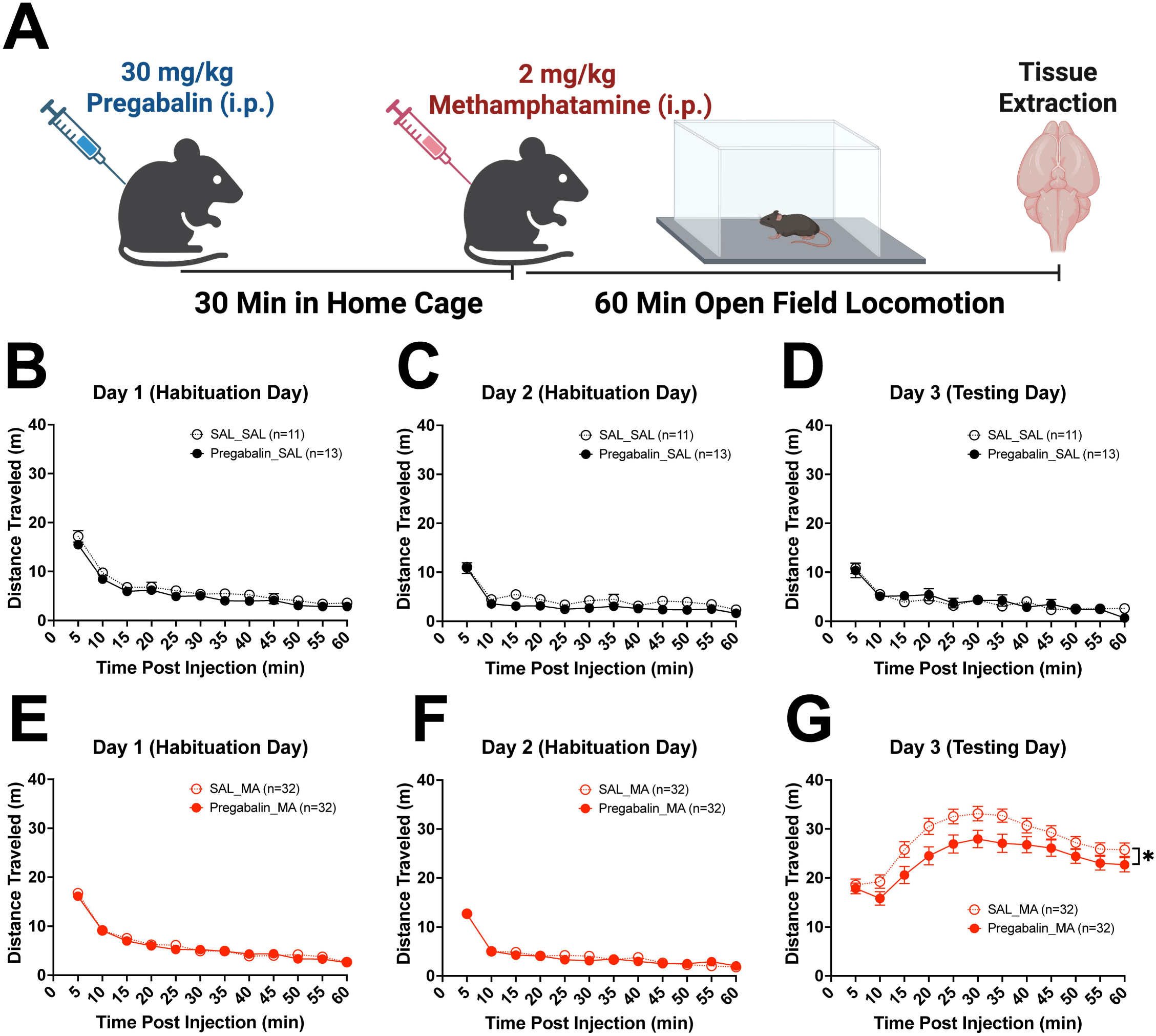
The CACNA2D2/CACNA2D1 inhibitor pregabalin selectively reduces methamphetamine-induced locomotor activity. On Days 1 and 2 (habituation days), 30 min after pretreatment with saline (i.p.) and placement back into the home cage, mice were injected again with saline (i.p.) and placed into testing apparatus for 1 h. On Day 3, approximately one-half of the mice were pretreated first with saline (i.p.) while the other half received pregabalin (30 mg/kg, i.p.) and were placed back into the home cage for 30 min. Mice were then injected with 2 mg/kg methamphetamine (i.p.) or saline (i.p.) and immediately placed into the testing apparatus for 60 min followed by immediate sacrifice and removal of the striatum. SAL_SAL = saline pre-treatment followed by saline; Pregabalin_SAL = pregabalin pre-treatment followed by saline; SAL_MA = SAL pre-treatment followed by MA; Pregabalin_MA = pregabalin pre-treatment followed by. **(A):** Injection timeline for those mice pre-treated with pregabalin (30 mg/kg, i.p.) and methamphetamine (2 mg/kg, i.p.) on Day 3. **(B-C):** Locomotor activity on Day 1 **(B)** and day 2 **(C)** for 1 hr in 5-min bins. Mice assigned for eventual pretreatment with saline or pregabalin followed by saline on Day 3 (left) showed no difference in locomotor activity on Day 1 [F(11,22)_Pregabalin_treatment_ = 0.793, p = 0.194; F(11,242)_Pregabalin_treatment x Time_ = 0.269, p = 0.990] and Day 2 [F(11,22)_Pregabalin_treatment_ = 2.204, p = 0.103; F(11,242)_Pregabalin_treatment x Time_ = 0.653, p = 0.782], indicating no spurious effect of random treatment assignment. **(D):** Pregabalin pre-treatment did not alter SAL-induced locomotor activity on Day 3 [F(11,22)_Pregabalin_treatment_ = 0.005, p = 0.946; F(11,242)_Pregabalin_treatment x Time_ = 1.485, p = 0.137]. **(E-F):** Locomotor activity on Day 1 **(E)** and Day 2 **(F)** for 1 hr in 5-min bins. Mice assigned for eventual pretreatment with saline or pregabalin followed by methamphetamine on Day 3 (left) showed no difference in locomotor activity on Day 1 [F(11,62)_Pregabalin_treatment_ = 0.238, p = 0.627; F(11,682)_Pregabalin_treatment x Time_ = 0.680, p = 0.758] and Day 2 [F(11,62)_Pregabalin_treatment_ = 0.126, p = 0.724; F(11,682)_Pregabalin_treatment x Time_ = 1.314, p = 0.332], again indicating no spurious effect of random treatment assignment. **(G):** Pregabalin pre-treatment significantly decreased methamphetamine-induced locomotor activity [F(11,62)_Pregabalin_treatment_ = 4.267, p = 0.043; F(11,683)_Pregabalin_treatment x time_ = 2.515, p = 0.004]. Sample size of each group is indicated in the legend. Data are presented as the mean ± S.E.M.

Because CLIP and transcriptome analysis suggested differential 3’UTR usage of *Cacna2d2* in response to MA, we examined 3’UTR usage in whole striatal tissue in response to pregabalin in saline- and methamphetamine-treated animals. Because pregabalin inhibits CACNA2D2 and CACNA2D1 and because *Cacna2d2* is an hnRNP H target, we wanted to know if pregabalin pre-treatment would lead to a cellular adaptive response as measured via changes in *Cacna2d2* versus *Cacna2d1* transcripts under saline and methamphetamine conditions. We hypothesized that because hnRNP H targets the 3’ UTR of *Cacna2d2*, then pregabalin could affect 3’UTR usage of *Cacna2d2*. We designed primers to test for differential 3’UTR usage of *Cacna2d2* via qPCR (**Figure 7A**), with primers targeting the distal vs. proximal 3’UTR, which are separated by a polyadenylation site and therefore could indicate differential 3’UTR usage between conditions. Additionally, we used existing primers that target *Cacna2d1* at the final exon junction just proximal to the 3’UTR as well (González *et al*., 2017). There was no effect of pregabalin on proximal 3’UTR *Cacna2d2* usage or distal 3’UTR usage between genotypes in mice administered PGB followed by saline (**Figures 7B-C)**. Additionally, there were no differences in *Cacna2d1* levels between PGB conditions in these same mice **(Figure 7D)**. Following methamphetamine, pregabalin induced an increase in proximal 3’UTR usage of *Cacna2d2* that just missed statistical significance (p = 0.051; **Figure 7E**). Intriguingly, there was no effect of pregabalin on *Cacna2d2* distal 3’UTR usage or on *Canca2d1* transcript levels in either genotype following methamphetamine (**Figure 7F-G**). Overall, our results provide evidence that under methamphetamine treatment, pharmacological inhibition of CACNA2D2 could lead to increased proximal 3’UTR usage of *Cacna2d2* which could ultimately affect CACNA2D2 protein levels, cellular responses (e.g., synaptic transmission), and behavior.

**Figure 7.**
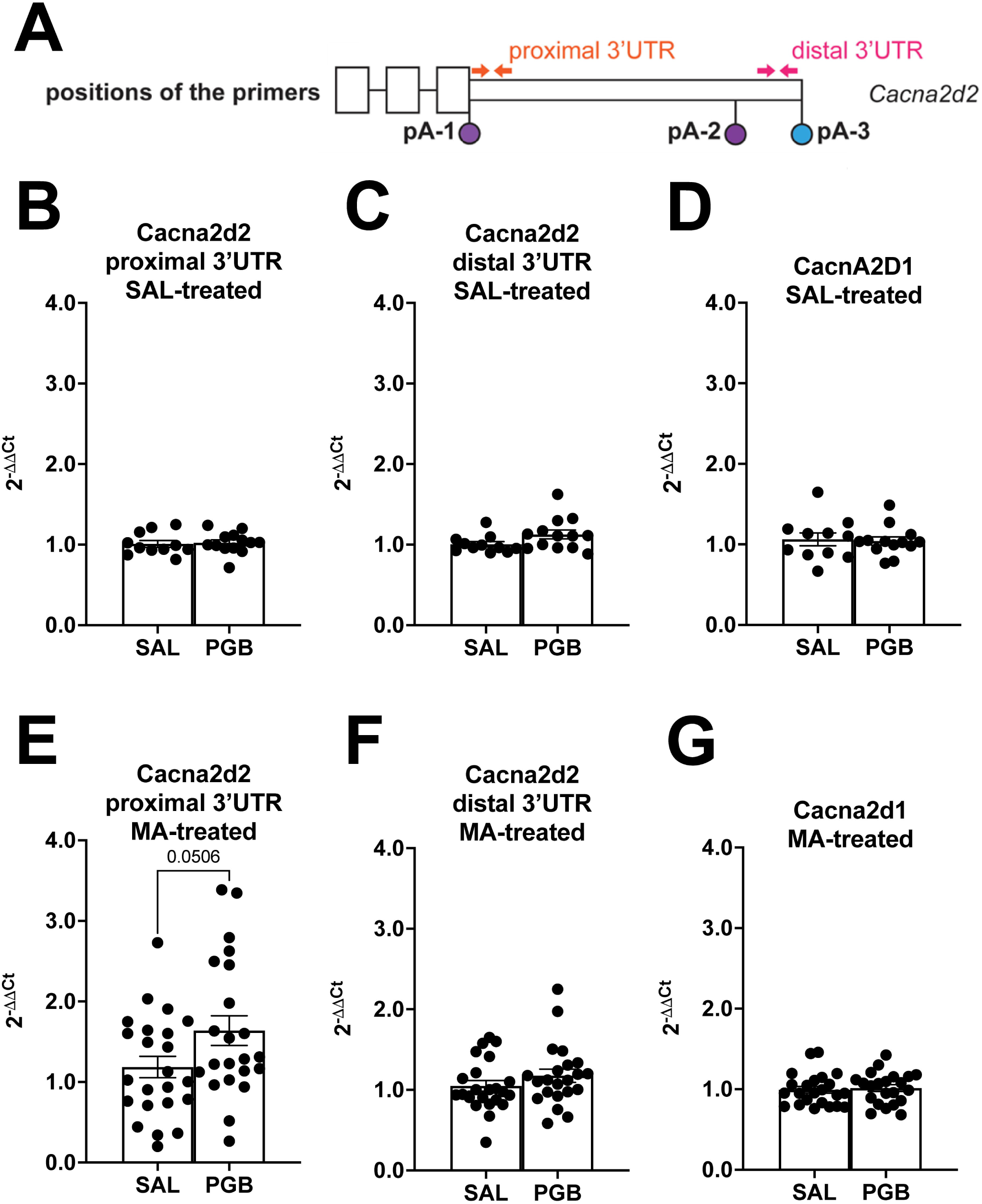
Pregabalin pretreatment increases proximal 3’ UTR usage of Cacna2d2 in striatum. 60 min post-saline or methamphetamine injection (90 min post-saline or pregabalin pretreatment), mice from the locomotor study (Figure 6) were sacrificed and whole striatum for each mouse was dissected and harvested for RT-qPCR to detect differential 3’UTR usage of Cacna2d2. SAL = saline; MA = methamphetamine; PGB = pregabalin. **(A):** Three polyadenylation sites (pA-1, pA-2, and pA-3) are present within the 3’UTR of *Cacna2d2* that distinguish isoforms containing 3’UTR of different lengths (UCSC Genome Browser). Primers were designed to detect differences in usage of the proximal and distal end of the 3’UTR of Cacna2d2 (UCSB annotations). The schematic indicates the positions of these primers. **(B-D):** In examining the effect of pregabalin pretreatment (30 mg/kg, i.p.) followed by saline (i.p.), there was no difference in the usage of the proximal 3’UTR of Cacna2d2 **(B)** [t(22) < 1, p = 0.811], no difference in the usage of the distal 3’UTR of Cacna2d2 **(C)** [t(22) = 1.791, p = 0.087], and no difference in Cacna2d1 transcript levels **(D)** [t(22) < 1, p = 0.819]. **(E-G):** Pre-treatment with pregabalin (30 mg/kg, i.p.) increased the proximal 3’UTR of Cacna2d2 in response methamphetamine (2 mg/kg, i.p.) relative to saline **(E)** [t(43) = 2.011, p = 0.051], without changing distal 3’UTR of Cacna2d2 **(F)** [t(43) = 1.195, p = 0.239], or Cacna2d1 transcript levels [t(43) = 0.363, p = 0.718]. Data are presented as the mean ± S.E.M.

## 4. DISCUSSION

This study defined the basal and methamphetamine-induced hnRNP H striatal targetome *ex vivo* from C57BL/6J wild-type mice which showed rapid methamphetamine-induced plasticity and provided a unique window into methamphetamine-induced gene regulation by a behaviorally relevant RNA-binding protein. hnRNP H localizes primarily to the nucleus of neurons in adult mice (Kamma et al., 1995; van Dusen et al., 2010; Ruan et al., 2020a), with low levels in cytoplasm (Wall et al., 2020). Nuclear binding of RBPs to 3’UTRs guides cytoplasmic localization and translation of mRNAs (Guramrit et al., 2015). Thus, nuclear hnRNP H binding to 3’UTRs of mRNAs could recruit other proteins necessary for export of mRNAs to cytoplasm for translation. Accordingly, most striatal hnRNP H targets were enriched for synaptic transmission (**Figure 2**). RBP-bound mRNAs are often functionally related because they share sequence elements recognized by the RBP and are post-transcriptionally regulated in concert (Keene and Tenenbaum, 2002). For hnRNP H, the core motifs in most target mRNAs were G-rich sequences (**Figure 1**), consistent with prior reports (Russo et al., 2010; Lefave et al., 2011; Huelga et al., 2012; Uren et al., 2016).

We observed multiple genes that showed changes in binding to hnRNP H, expression, and exon usage in response to methamphetamine, with a small list of genes showing multiple types of regulation (**Figure 4**). One of these genes was *Kalrn,* in which methamphetamine treatment reduced hnRNP H binding to intron 1 of this gene. *Kalrn* codes for the KALIRIN protein in mice, a dendritically localized synaptic guanine nucleotide exchange factor for Rho-GTPases whose dysregulation is linked to various neurogenerative and neurodevelopmental conditions like autism and Alzheimer’s disease (Paskus *et al*., 2020; Parnell *et al*., 2021). Importantly, KALIRIN expression has been linked extensively with cocaine outcomes in rodents, with knockout of the KALIRIN-7 isoform specifically causing increase cocaine self-administration in mice (Kiraly *et al*., 2013), while in rats repeated cocaine injections increased KALIRIN-7 expression, along with KALIRIN-7 overexpression associated with an increased AMPA currents from medium spiny neurons in the nucleus accumbens (Ma *et al*., 2012; Wang *et al*., 2013). Interestingly, daily cocaine injections induced *Kalrn* promotor usage in the nucleus accumbens, potentially contributing to synaptic transmission and behavior (Mains *et al*., 2011). Therefore, hnRNP H-dependent *Kalrn* splicing could mediate methamphetamine neurobehavioral effects. Only one other gene that exhibited differential hnRNP H binding and differential exon usage, *Arhgef26,* had previously been associated with psychostimulant behavior, with Arhgef26 transcript levels in nucleus accumbens associated with cocaine infusions in mice in a genome-wide association study (Khan *et al*., 2023). However, multiple other genes that met these criteria were also implicated in neurodevelopmental disorders. These include *Dctn1,* important for axonal transport and linked to the neurodevelopmental disease Perry Syndrome plus frontotemporal dementia and amyotrophic lateral sclerosis (Konno *et al*., 2017; Dulski *et al*., 2010), *Ubqln1*, which is important for cell death and whose overexpression is associated with Alzheimer’s Disease (Jantrapirom *et al*., 2019), and *Agrn,* whose mutation leads to motor neuron death and is associated with subsequent muscle weakness (Jacquier *et al*., 2022). As multiple molecular alterations that underlie multiple neurodegenerative diseases, including mitochondrial alterations and oxidative stress, are also induced through substance misuse (including psychostimulants like cocaine and methamphetamine; Guo *et al*., 2023), these genes would be useful to follow up on and could plausibly link hnRNP H binding to methamphetamine locomotor increases.

*Mid1* was a top methamphetamine-upregulated gene that also exhibited differential binding to hnRNP H. *Mid1* is a microtubule-associated gene that contributes to axon developmental regulation (Lu *et al*., 2013). *Mid1* is particularly important for cerebellar neuromotor development (Dierssen *et al*., 2012; Nakamura *et al*., 2017), and while the gene has no known role in addiction, mutations have been shown to influence the development of cancer and neurodegenerative diseases (Krauss, 2016). Finally, while not a differential hnRNP H target, *Piezo1* showed significant gene expression and differential exon usage in response to methamphetamine. *Piezo1* codes for a mechanosensitive ion channel important for central nervous system physiology, and is implicated in multiple central nervous system disorders including Alzheimer’s, stroke, and neuroinflammation (Xu *et al*., 2024). To summarize, multiple genes associated with neurodegeneration appear to be regulated by methamphetamine treatment and in some cases, linked to hnRNP H binding.

*Cacna2d2* emerged as an interesting gene as it showed evidence for differential binding, 3’UTR usage, and gene expression in response to methamphetamine, suggesting that its rapid, dynamic regulation could mediate methamphetamine-induced locomotor behavior. In support, pharmacological inhibition of CACNA2D2 with pregabalin decreased methamphetamine-induced locomotor activity. Further support came from the observation that pregabalin pretreatment followed by methamphetamine administration induced a near-significant striatal upregulation of Cacna2d2 transcripts containing the proximal 3’ UTR (**Figure 7**), whereas there was no effect on Cacna2d1 transcripts. 3’UTR alt-splicing results in different CACNA2D2 protein isoforms created (Barclay and Rees, 2000). Interestingly, the most abundant RNA isoform contains only the proximal 3’UTR and not the distal. Therefore, it’s possible that differential hnRNP H1 binding at the 3’UTR may ultimately lead to differential CACNA2D2 protein isoforms between genotypes, influencing how pregabalin binds and exerts its effects. Pregabalin binds to both CACNA2D2 and CACNA2D1 subunits with similar affinities and in several brain regions, including striatum (Li et al., 2011). As both CACNA2D2 and CACNA2D1 subunits are expressed in the rodent striatum (Barclay and Rees, 2000; Taylor and Garrido, 2008), its plausible that pregabalin binding to either CACNA2D2 and/or CACNA2D1 could lower methamphetamine-induced locomotion. However, the negative finding regarding pregabalin and Cacna2d1 expression suggests that the behavioral effects of pregabalin could be mediated specifically by CACNA2D2. Interestingly, the hnRNP H RNA-binding targets containing the most common motif (G-rich) in their binding sites were highly enriched for “presynaptic depolarization and calcium channel opening” pathway (**Figure 1D**), further implicating altered calcium activity in linking hnRNP H activity to methamphetamine locomotion.

*Cacna2d2* codes for a pre-protein that is processed into α2 and δ2 subunits of voltage-gated calcium channels (**VGCCs**) (Dolphin and Lee, 2020) and is expressed in cerebellum, striatum, and hippocampus (Dolphin, 2012). α2δ2 subunits localize VGCCs to the active zone and promote neurotransmitter release (Dolphin, 2013). The effects of α2δ2 subunits on synaptic transmission are complex. Overexpression of α2δ2 subunits decreased presynaptic calcium elevation following an action potential, yet increased vesicular release (Hoppa et al., 2012). Additionally, overexpression of Cacna2d2 disrupted intracellular calcium signaling and mitochondrial function (Carboni et al., 2003). Following calcium influx, mitochondria can buffer intracellular calcium (Rizzuto et al., 2012). Our previous proteomic analysis of the striatal synaptosome showed methamphetamine-induced changes in synaptic mitochondrial protein levels (Ruan et al., 2020a). Methamphetamine inhibits calcium entry into L-type and N-type VGCCs and longer exposure induces a compensatory upregulation of *Cacna1c* transcript in SH-SY5Y cells (Andres et al., 2015). Intracellular and extracellular calcium contribute to methamphetamine-induced dopamine release in the ventral striatum (Yorgason et al., 2020). The rapid, methamphetamine-induced regulation of Cacna2d2 could reflect an adaptive cellular response to control calcium entry and synaptic transmission.

Pregabalin is used to treat seizures (Panebianco et al., 2019), neuropathic pain (Attal et al., 2010), and anxiety (Slee et al., 2019). Our findings are the first to show that pregabalin can attenuate methamphetamine-induced locomotor stimulation and compliment studies showing pregabalin-induced decreases in ketamine-(Nunes et al., 2012), morphine-induced locomotion (Vashchinkina et al., 2018), and in cocaine oral self-administration (de Guglielmo et al., 2013). Prior work with gabapentin, primarily a CACNA2D1 inhibitor but with some affinity for CACNA2D2, showed a reduction in methamphetamine-induced locomotor sensitization (Kurokawa et al., 2010). Two recent studies found that the most commonly co-misused substance type with illicit pregabalin misuse was amphetamine-like compounds, including methamphetamine (Ovat *et al*., 2024; Sağlam *et al*., 2025). Although this observation raises several questions as to why this is the case, one intriguing possibility is that users of amphetamines are in some way self-medicating to stave off the aversive effects of psychostimulant withdrawal and to normalize their physiological state. This hypothesis could be tested at the preclinical level via in vivo or ex vivo electrophysiology.

To assess the effect of pregabalin on Cacna2d2 expression, we quantified Cacna2d2 mRNA in tissue harvested 60 min post-methamphetamine injection and 90 min post-pregabalin injection (**Figure 7**). This was a strategic decision upfront to ensure we would not miss any time window where pregabalin might have an effect. In retrospect, pregabalin’s behavioral effects were most pronounced within the first 30 min; thus, analyzing *Cacna2d2* expression at 30 min post-methamphetamine might yield more robust results. Furthermore, a limitation of our design was that we could not directly compare methamphetamine treatment effects on *Cacna2d2* expression. Our rationale for conducting separate experiments for methamphetamine and saline was that we first wanted to determine if there was an effect of pregabalin on methamphetamine behavior. Upon observing a significant behavioral result, we then ran the experiment involving saline-treated control mice, to rule out non-specific effects of pregabalin on locomotion. This sequential strategy would have prevented us from using an excess number of mice, had we obtained a null result of pregabalin on methamphetamine behavior.

To conclude, we provide the first methamphetamine-induced RNA targetome on an RBP, hnRNP H. Analysis of drug-induced RBP targetomes, especially drugs of abuse, is a novel, understudied approach for understanding rapid, synaptic gene regulation as it relates to cell biological adaptations in neuronal excitability, neurotransmitter release, plasticity, and behavior. We focused on hnRNP H, given the evidence for its role in methamphetamine-induced dopamine release and addiction model behaviors (Yazdani et al., 2015; Bryant and Yazdani, 2016; Ruan et al., 2020a, 2020b) across multiple substances (Bryant et al., 2020; Fultz et al., 2021).

We established hnRNP H as a novel gene regulatory link among several target mRNAs coding for proteins known to mediate psychostimulant-induced neurotransmission and plasticity. Finally, we pharmacologically validated CACNA2D2 as a functional mechanistic target linking hnRNP H with methamphetamine behavior. Our study design represents a powerful approach for triangulating complementary-omics datasets that can be broadly applied to the study of drug-induced RBP-RNA dynamics and discovery of RNA-binding targets underlying cell biological responses and neurobehavioral adaptations that can be leveraged for novel therapeutics.

## Declaration of competing interest

The authors declare no competing financial or personal interests.

## Acknowledgement

This work was supported by the National Institutes of Health – National Institute on Drug Abuse [grant numbers R01DA039168, U01DA050243; F31DA056217, T32DA055553]; and National Institutes of Health – National Institute of General Medical Sciences [grant number R25GM125511]; and the Boston University Undergraduate Research Program.

## CRediT authorship contribution statement

**Qiu T. Ruan:** Conceptualization, Data curation, Formal analysis, Investigation, Methodology, Supervision, Validation, Visualization, Writing – original draft, Writing – review and editing. **William B. Lynch:** Conceptualization, Data curation, Formal analysis, Investigation, Methodology, Supervision, Validation, Visualization, Writing – original draft, Writing – review and editing. **Rebecca H. Cole:** Formal analysis, Investigation**. Michael A. Rieger:** Investigation, Supervision. **Britahny M. Baskin:** Investigation. **Sophia A. Miracle:** Investigation. **Jacob A. Beierle:** Investigation. **Emily J. Yao:** Investigation. **Jiayi W. Cox:** Investigation. **Amarpreet Kandola:** Investigation. **Kayla T. Richardson:** Investigation. **Melanie M. Chen:** Investigation. **Julia C. Kelliher:** Data Curation, Investigation. **R. Keith Babbs:** Formal analysis, investigation. **Peter E. A. Ash:** Resources, Supervision. **Benjamin Wolozin:** Resources. **Karen K. Szumlinski:** Resources. **W. Evan Johnson:** Methodology, Resources, Software. **Joseph D. Dougherty:** Resources. **Camron D. Bryant:** Conceptualization, Funding acquisition, Project administration, Resources, Supervision, Writing – review and editing.

## Supplementary Information

**Figure S1. Optimization of the CLIP conditions for hnRNP H. (A):** Immunoblot shows the IP condition using the anti-hnRNP H antibody. No nonspecific bands were detected in the rabbit IgG pulldown. In the hnRNP H IP, a band of 50 kDa corresponding to the size of hnRNP H was detected. **(B):** Immunoblot showing IP condition using different concentrations of hnRNP H antibody. A total of 20 ug of antibody was chosen for the IP. **(C):** RNA ^32^P autoradiogram showing bound RNA under different duration of RNase I_f_ digestion. Three min was chosen as the optimal length of digestion.

**Figure S2. cDNA libraries generation followed by visualization with DNA gel electrophoresis.** (A): No cDNA library was generated from IgG mock IP. Even after 28 cycles, no cDNA library (which should be > 200 bp) was detected. For this reason, these four samples were not subjected to RNA-seq. (B): In contrast with CLIP cDNA libraries, DNA bands > 200 bp were detected after 20 PCR cycles.

**Table S1. Read Coverage in CLIP-seq and input RNA-seq samples.**

**Table S2. CLIP-seq identified hnRNP H peaks for saline treated animals.**

**Table S3. CLIP-seq identified hnRNP H peaks for methamphetamine treated animals.**

**Table S4. hnRNP H motif by genomic region in saline treatment.**

**Table S5. Top 10 pathways enriched in hnRNP H-associated targets with G-rich motif in saline treatment.**

**Table S6. hnRNP H binding sites on the 7 targets enriched for “presynaptic depolarization and calcium channel opening**.”

**Table S7. hnRNP H RNA binding targets between methamphetamine versus saline.**

**Table S8. Chi-square tests comparing difference in proportion of hnRNP H associated binding regions.**

**Table S9. GO pathway enrichment for hnRNP H RNA binding targets between methamphetamine versus saline.**

**Table S10. Differential expressed genes between methamphetamine versus saline.**

**Table S11. Genes with differential exon and intron usage between methamphetamine versus saline.**

## Notes

https://www.ncbi.nlm.nih.gov/geo/query/acc.cgi?acc=GSE160682

### Competing Interest Statement

The authors have declared no competing interest.

### Summary of Updates

We have removed certain data due to recent discoveries as detailed in the manuscript. We have also re-run the analyses and provided updated figures and statistics with the current data. Findings from the current data are interpreted and discussed in the updated manuscript.

