## Supplementary figures and images for "The striatal heterogeneous nuclear ribonucleoprotein H mRNA targetome associated with methamphetamine administration and behavior"

### Supplemental Figure 1

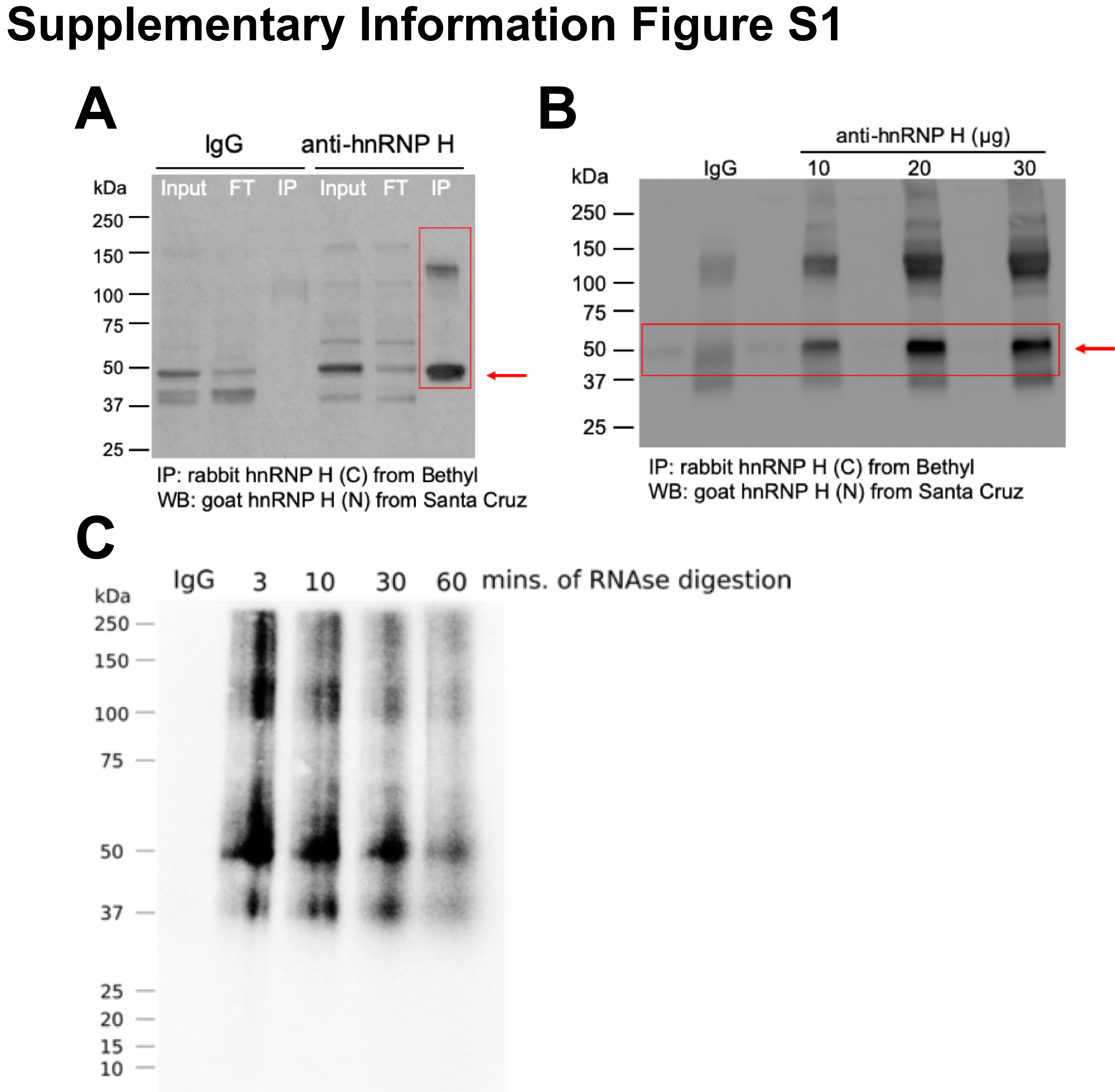

### Supplemental Figure 2

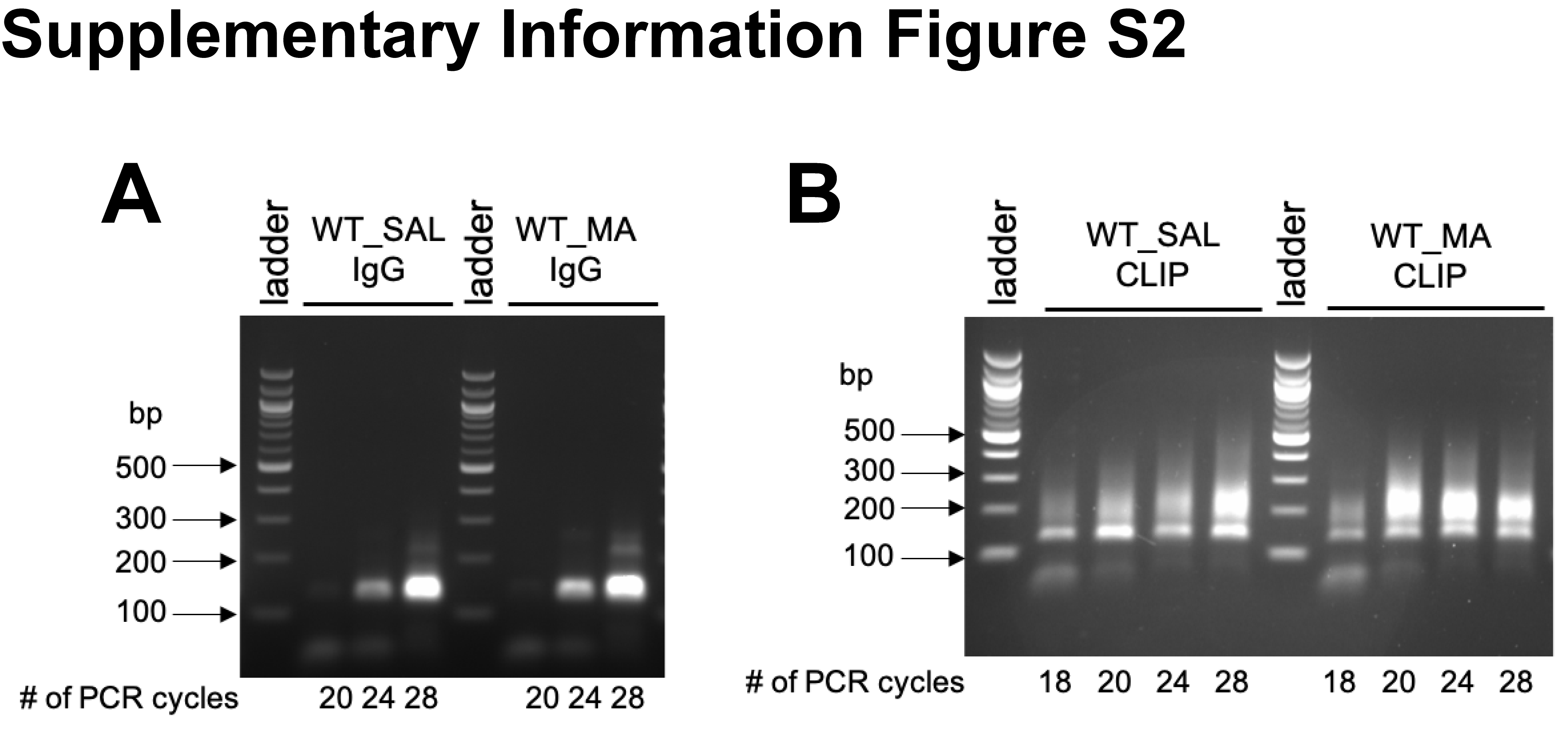
