## Supplemental Tables 1, 4, 5, 6, 8 for "The striatal heterogeneous nuclear ribonucleoprotein H mRNA targetome associated with methamphetamine administration and behavior"

**Table S1. Read Coverage in CLIP-seq and input RNA-seq samples.**

| **Sample Name** | **Type** | **# of Mice** | **Genotype** | **Treatment** | **Replicate** | **Barcode Sequence** | **Read Counts** |
| --- | --- | --- | --- | --- | --- | --- | --- |
| CLIP_WT_SAL_1 | CLIP-seq | 2 M + 2 F | WT | SAL | 1 | ACAGTG | 16,446,905 |
| CLIP_WT_SAL_2 | CLIP-seq | 2 M + 2 F | WT | SAL | 2 | GATCAG | 12,440,933 |
| CLIP_WT_SAL_3 | CLIP-seq | 2 M + 2 F | WT | SAL | 3 | AGTCAA | 22,826,085 |
| CLIP_WT_MA_1 | CLIP-seq | 2 M + 2 F | WT | MA | 1 | GCCAAT | 24,419,809 |
| CLIP_WT_MA_2 | CLIP-seq | 2 M + 2 F | WT | MA | 2 | TAGCTT | 16,402,702 |
| CLIP_WT_MA_3 | CLIP-seq | 2 M + 2 F | WT | MA | 3 | AGTTCC | 13,376,067 |
| INPUT_WT_SAL_1 | RNA-seq | 2 M + 2 F | WT | SAL | 1 | GTTTCG | 44,421,956 |
| INPUT_WT_SAL_2 | RNA-seq | 2 M + 2 F | WT | SAL | 2 | ACTGAT | 48,930,085 |
| INPUT_WT_SAL_3 | RNA-seq | 2 M + 2 F | WT | SAL | 3 | CAACTA | 53,470,359 |
| INPUT_WT_MA_1 | RNA-seq | 2 M + 2 F | WT | MA | 1 | CGTACG | 53,843,037 |
| INPUT_WT_MA_2 | RNA-seq | 2 M + 2 F | WT | MA | 2 | ATGAGC | 55,822,205 |
| INPUT_WT_MA_3 | RNA-seq | 2 M + 2 F | WT | MA | 3 | CACGAT | 44,088,258 |

**Table S4. hnRNP H motif by genomic region in saline treatment.** The top 3 Homer de novo motif results are shown for each genomic region type (5’UTR, CDS, intron and 3’UTR). The motif discovery was performed in Homer (Heinz et al., 2010).

| **Genomic Region Type** | **Motif** | **p value** | **% of Target** |
| --- | --- | --- | --- |
| **5’UTR** | 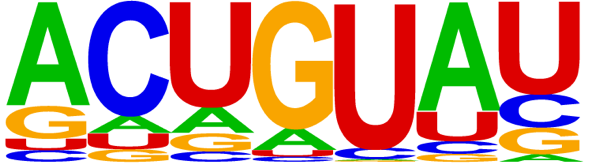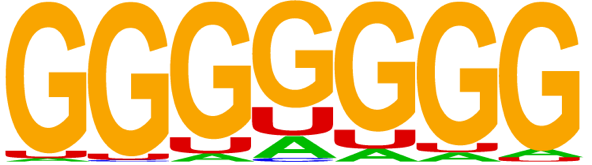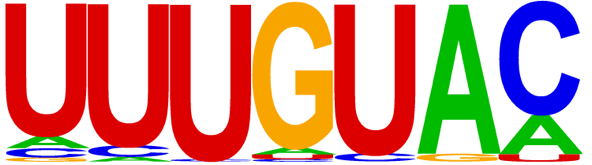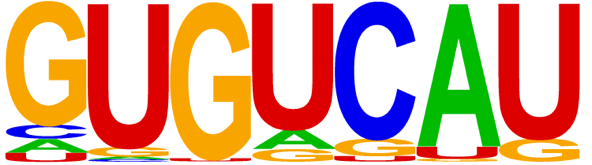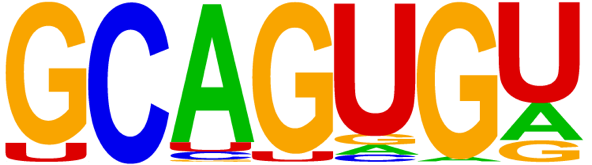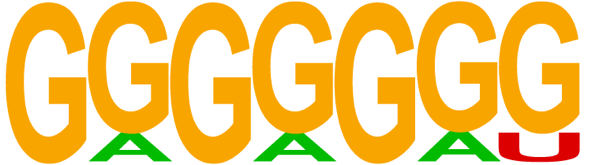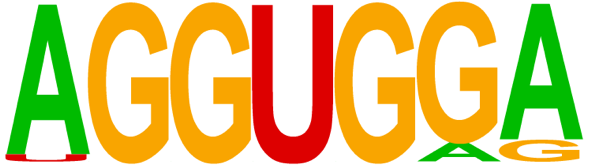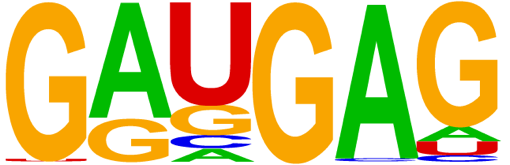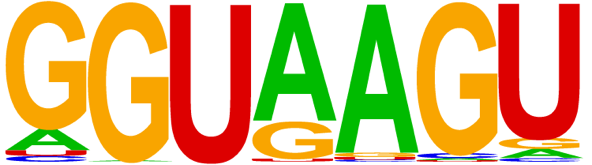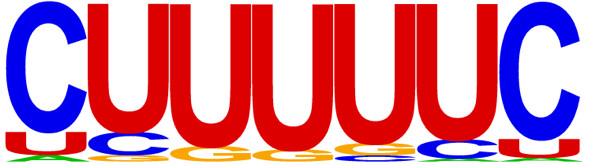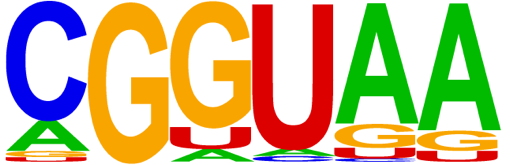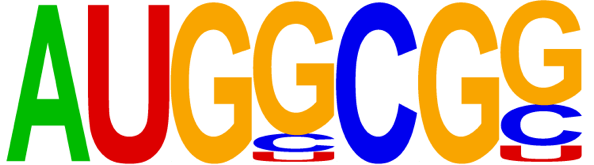 | 1e-20 | 5.57% |
|  |  | 1e-16 | 23.49% |
|  |  | 1e-14 | 10.87% |
| **CDS** |  | 1e-122 | 18.76% |
|  |  | 1e-75 | 15.00% |
|  |  | 1e-70 | 18.36% |
| **Intron** |  | 1e-47 | 12.19% |
|  |  | 1e-34 | 9.88% |
|  |  | 1e-34 | 7.75% |
| **3’UTR** |  | 1e-54 | 15.74% |
|  |  | 1e-47 | 9.85% |
|  |  | 1e-42 | 30.08% |

**Table S5. Top 10 pathways enriched in hnRNP H-associated targets with G-rich motif in saline treatment.**

| **Pathway** | **Database** | **Adjusted p** | **RNA targets** |
| --- | --- | --- | --- |
| Presynaptic depolarization and calcium channel opening | pathway_Reactome | 9.52E-05 | *Cacna1a, Cacnb4, Cacna1e, Cacng4, Cacna1b, Cacng2, Cacnb2* |
| Phase 1 - inactivation of fast Na+ channels | pathway_Reactome | 0.0225 | *Kcnd3, Kcnip3, Kcnip2, Kcnd2* |
| LGI-ADAM interactions | pathway_Reactome | 6.95E-04 | *Dlg4, Adam23, Stx1b, Lgi4, Cacng4, Cacng2, Adam22* |
| Unblocking of NMDA receptors, glutamate binding and activation | pathway_Reactome | 1.85E-04 | *Gria1, Dlg4, Dlg2, Grin2a, Gria3, Camk2d, Grin2b, Actn2, Camk2a* |
| Interaction between L1 and Ankyrins | pathway_Reactome | 0.0186 | *Sptbn4, Ank3, Ank2, Ank1, Sptb* |
| CREB phosphorylation through the activation of CaMKII | pathway_Reactome | 2.86E-04 | *Dlg4, Creb1, Dlg2, Grin2a, Calm1, Camk2d, Grin2b, Actn2, Camk2a* |
| Ras activation upon Ca2+ influx through NMDA receptor | pathway_Reactome | 4.41E-04 | *Rasgrf1, Dlg4, Dlg2, Grin2a, Calm1, Camk2d, Grin2b, Actn2, Camk2a* |
| Reduction of cytosolic Ca++ levels | pathway_Reactome | 0.0096 | *Atp2a2, Atp2b2, Slc8a2, Atp2b1, Calm1, Slc8a1* |
| Rap1 signaling | pathway_Reactome | 0.0096 | *Rap1gap, Ywhaz, Prkg1, Rasgrp2, Rap1gap2, Rasgrp1* |
| Hypothetical Network for Drug Addiction | pathway_Wikipathway | 1.76E-05 | *Cacna1a, Cacnb4, Cacna1e, Cacng4, Cacna1b, Cacng2, Cacnb2* |

**Table S6. hnRNP H binding sites on the 7 targets enriched for “presynaptic depolarization and calcium channel opening**.”

| **RNA-binding Target** | **Peak Position(s)** | **Genomic Region Type** |
| --- | --- | --- |
| *Cacna1a* | chr8: 84611302-84611402 | Intron |
| *Cacnb4* | chr8: 52629320- 52629420  chr8: 52560620- 52560720  chr8: 52472520- 52472620 | Intron  Intron  Intron |
| *Cacna1e* | chr1:154446931- 154447031 | Intron |
| *Cacng4* | chr11:107794366- 107794466 | 5’UTR or CDS depending on the mRNA isoform |
| *Cacna1b* | chr2: 24608087- 24608187  chr2: 24606587-24606687 | CDS or intron depending on the mRNA isoform  CDS |
| *Cacng2* | chr15: 78045548 - 78045548  chr15: 78103948- 78104048 | Intron  Intron |
| *Cacnb2** | Chr2: 14762588- 14762688  Chr2:14685188- 14685288  Chr2: 14622788- 14622888 | Intron  Intron  Intron |

***only 3 out of 9 peaks lists**

**Table S8. Chi-square tests comparing difference in proportion of hnRNP H associated binding regions.** Given the binding events associated with hnRNP H detected in in saline (SAL) treatment, the proportions of 3’UTR and intron targets in methamphetamine (MA) treatment are significantly differently as depicted by the p values calculated in chi square test.

| Subregion | MA vs SAL |
| --- | --- |
| 3'UTR | 0.0008 (***) |
| distal intron | < 0.0001 (****) |
| proximalx500_intron | < 0.0001 (****) |
| proximax200_intron | < 0.0001 (****) |
| other_exon | 0.42 |
| CDS | 0.208 |
| 5'UTR | 0.237 |
